# Identification of hyperosmotic stress-responsive genes in Chinese hamster ovary cells via genome-wide virus-free CRISPR/Cas9 screening

**DOI:** 10.1101/2022.12.13.520335

**Authors:** Su Hyun Kim, Seunghyeon Shin, Minhye Baek, Kai Xiong, Karen Julie la Cour Karottki, Hooman Hefzi, Lise Marie Grav, Lasse Ebdrup Pedersen, Helene Faustrup Kildegaard, Nathan E. Lewis, Jae Seong Lee, Gyun Min Lee

**Affiliations:** Department of Biological Sciences, KAIST, Daejeon, 34141, Republic of Korea; The Novo Nordisk Foundation Center for Biosustainability, Technical University of Denmark, Lyngby, Denmark; Departments of Pediatrics and Bioengineering, University of California, San Diego, USA; Department of Molecular Science and Technology, Ajou University, Suwon, 16499, Republic of Korea

**Keywords:** Chinese hamster ovary cell, CRISPR/Cas9 screen, Genome-wide screen, Osmotic stress, Therapeutic protein

## Abstract

Chinese hamster ovary (CHO) cells are the preferred mammalian host cells for therapeutic protein production that have been extensively engineered to possess the desired attributes for high-yield protein production. However, empirical approaches for identifying novel engineering targets are laborious and time-consuming. Here, we established a genome-wide CRISPR/Cas9 screening platform for CHO-K1 cells with 111,651 guide RNAs (gRNAs) targeting 21,585 genes using a virus-free recombinase-mediated cassette exchange-based gRNA integration method. Using this platform, we performed a positive selection screening under hyperosmotic stress conditions and identified 180 genes whose perturbations conferred resistance to hyperosmotic stress in CHO cells. Functional enrichment analysis identified hyperosmotic stress responsive gene clusters, such as tRNA wobble uridine modification and signaling pathways associated with cell cycle arrest. Furthermore, we validated 32 top-scoring candidates and observed a high rate of hit confirmation, demonstrating the potential of the screening platform. Knockout of the novel target genes, *Zfr* and *Pnp*, in monoclonal antibody (mAb)-producing recombinant CHO (rCHO) cells and bispecific antibody (bsAb)-producing rCHO cells enhanced their resistance to hyperosmotic stress, thereby improving mAb and bsAb production. Overall, the collective findings demonstrate the value of the screening platform as a powerful tool to investigate the functions of genes associated with hyperosmotic stress and to discover novel targets for rational cell engineering on a genome-wide scale in CHO cells.

## 1. Introduction

Chinese hamster ovary (CHO) cells have been widely used for the large-scale production of recombinant biotherapeutics (Walsh, 2018; Wurm, 2013). Extensive efforts have been made in CHO cell engineering to achieve high-yield production with improved product quality and low manufacturing costs (Kim et al., 2012; Tihanyi and Nyitray, 2021). However, as current cell engineering strategies mostly rely on known targets, new strategies are required to identify novel targets in CHO cells.

Clustered regularly interspaced short palindromic repeats (CRISPR)/CRISPR-associated protein 9 (Cas9) technology has facilitated high-throughput large-scale functional screening, which can be used to identify novel targets associated with a specific phenotype of interest (Shalem et al., 2014; Wang et al., 2014). Being a powerful tool for gene perturbations, CRISPR screens have been widely used in various applications, such as investigation of molecular and cellular biology, genetic disease, cancer, and microbial engineering (Bock et al., 2022). Recently, a metabolic CRISPR/Cas9 screen in CHO cells identified a novel target, whose deletion improved cell growth in glutamine-free media (Karottki et al., 2021). Therefore, CRISPR genetic screening is a promising tool to discover novel targets for CHO cell engineering.

Most pooled CRISPR screens rely on lentivirus-based delivery methods for the introduction of guide RNAs (gRNAs) (Joung et al., 2017). However, working with lentiviruses requires advanced biosafety facilities, which may be cumbersome in terms of technical accessibility. Despite delicate adjustment of the multiplicity of infection, transducing viruses can cause no or multiple integrations of gRNA, following a Poisson distribution, which can decrease the signal-to-noise ratio (Ellis and Delbrück, 1939). To overcome these drawbacks, alternative approaches using plasmid transfection, such as Cas9-mediated homologous recombination (Rajagopal et al., 2016), recombinase-mediated cassette exchange (RMCE) (Viswanatha et al., 2018), and transposons (Chang et al., 2020) have provided new options to replace lentivirus-based methods. Notably, RMCE coupled with a landing pad platform, which encourages single gene integration into a pre-defined site, shows a higher efficiency of single-copy gRNA integration with minimized clonal variation compared to the lentivirus-based system (Xiong et al., 2021).

For the large-scale production of recombinant biotherapeutics, including monoclonal antibodies (mAbs), fed-batch culture supporting high volumetric productivity is widely used because of its operational simplicity and reliability (Fike, 2009). In fed-batch cultures, culture osmolality increases with culture time due to repeated feeding of nutrient concentrates and addition of a base to maintain optimal pH during culture, which induces apoptotic cell death (Han et al., 2010). Hence, the use of an apoptosis-resistant CHO cell line can further increase the volumetric productivity of recombinant biotherapeutics in fed-batch cultures by extending the culture duration. Recently, CRISPR/Cas9 screening using gRNA libraries targeting genes related to kinase and cell cycle identified an apoptosis-related target whose deletion conferred resistance to osmotic stress in rHEK293 cells (Shin et al., 2022). However, little is known about the functional characterization of genes related to hyperosmotic conditions in CHO cells on a genome-wide scale.

In this study, we performed genome-wide CRISPR knockout screening of CHO-K1 cells using a virus-free RMCE-based gRNA integration method to identify novel genes associated with hyperosmotic stress. A proliferation-based positive-selection screen was conducted against hyperosmotic stress conditions and genes that were targeted by significantly enriched gRNAs were identified. Functional enrichment analysis was conducted using the identified genes and hyperosmotic stress responsive gene networks were elucidated. Perturbations of the 32 highest-ranking genes, whose gRNAs showed significant enrichment on the screen, were validated in CHO-K1 cells. We focused on *Zfr* and *Pnp* genes and verified their perturbations in mAb- and bispecific antibody (bsAb)-producing recombinant CHO (rCHO) cell lines.

## 2. Materials and methods

### 2.1. Cloning and plasmid constructions

All plasmids used in this study are listed in Supplementary Table S1. A TagBFP RMCE donor plasmid and nuclear localization signal (NLS)-Bxb1 recombinase plasmid were constructed using the uracil-specific excision reagent (USER) cloning method, as previously described (Lee et al., 2015). TagBFP RMCE donor plasmid was generated from the attB-Puro-U6-gRNA-attB^mut^ plasmid (Xiong et al., 2021). TagBFP-coding sequence was amplified from the plasmid described previously (Sergeeva et al., 2020). NLS-Bxb1 recombinase plasmid was generated from the PSF-CMV-Bxb1 recombinase plasmid (Xiong et al., 2021). NLS sequences were included in the USER primers to be attached to both the N- and C-termini of Bxb1 recombinase. All primers used for cloning are listed in Supplementary Table S2. To generate an all-in-one CRISPR/Cas9 plasmid for screening verification, the annealed gRNA oligos were cloned into the BbsI site of the pSpCas9(BB)-2A-BSD plasmid (Addgene plasmid # 118055; a gift from Ken-Ichi Takemaru) using T4 ligase, according to the manufacturer’s instructions. All gRNAs used in this study are listed in Supplementary Table S3. All constructs were verified by sequencing and purified using a NucleoBond Xtra Midi EF kit (Macherey-Nagel, Düren, Germany), according to the manufacturer’s instructions.

### 2.2. Library design and construction

To construct a CHO-K1 genome-wide CRISPR knockout library, we designed 111,651 unique gRNAs against 21,585 genes. The library was designed and constructed as previously described (Xiong et al., 2021). DESKTOP Genetics (DESKGEN, London, UK) designed a whole-genome gRNA library against the Chinese hamster PICR scaffold. All protein-coding gene IDs were independently targeted. Ideally, five gRNAs were designed per gene that preferentially targeted the predicted functional domains. Additionally, 1000 non-targeting gRNAs were designed in the library as internal controls. This resulted in a total library of 108,580 gRNAs targeting 21,585 genes. To precisely target the CHO-K1 cell line, it was necessary to correct the library design. With a large number of Illumina reads from CHO samples at Denmark Technical University, it was possible to overlay CHO reads on the PICR scaffold and identify the mutations present in CHO-K1 cells. Mutations were categorized as indels and single nucleotide polymorphisms. If gRNAs bound to regions with mutations, the gRNA sequence was corrected to target the mutated sequence. This resulted in the selection of 3,071 gRNAs. For convenience, the corrected gRNAs for CHO-K1 genome were added directly to the library. Therefore, a final library consisting of 111,651 gRNAs targeting 21,585 genes was designed. The designed library was synthesized as an oligo by Twist Bioscience (San Francisco, CA).

### 2.3. Cell lines, culture maintenance, and culture media

CHO-K1 cells were cultured in Dulbecco’s modified Eagle’s medium (Gibco, Grand Island, NY) supplemented with 7% fetal bovine serum (HyClone, Logan, UT). Cells were maintained in T25 flasks at 37°C with 5% CO_2_ and passaged every three days. CHO-K1 cells were adapted to grow in suspension culture in 125 mL Erlenmeyer flasks (Corning, Corning, NY) containing 30 mL of CD CHO medium (Gibco) supplemented with 4 mM glutamine (HyClone) and 100X anti-clumping agent (Lonza, Basel, Switzerland) in a climo-shaking CO_2_ incubator (ISF1-X; Adolf Kuhner AG, Birsfelden, Switzerland) at 110 rpm, 37 °C, 5% CO_2_, and 85% humidity. CHO-K1 cell line producing rituximab (CHO-mAb) was established as previously described (Park et al., 2016). CHO-mAb cell line was maintained in PowerCHO2CD medium (Lonza) supplemented with glutamine synthetase expression medium (Sigma-Aldrich, St. Louis, MO), and 25 μM methionine sulfoximine (Sigma-Aldrich) in 125 mL Erlenmeyer flasks. CHO-S cell line producing bsAb (CHO-bsAb) was provided by ABL Bio (Gyeonggi-Do, Korea). CHO-bsAb cell line was maintained in Dynamis medium (Gibco) supplemented with 4 mM glutamine, 100 nM methotrexate (Sigma-Aldrich), and 0.2% anti-clumping agent (Thermo Fisher Scientific, Waltham, MA). Viable cells were distinguished from dead cells using the trypan blue dye exclusion method, and cell concentration was estimated using a Countess II FL automated cell counter (Invitrogen, Carlsbad, CA).

### 2.4. Generation of the landing pad master cell line (MCL) using the CRISPR/Cas9-based RMCE landing pad platform

CHO-K1 MCL was generated as previously described (Grav et al., 2018). CHO-K1 cells (0.5 × 10^6^ cells/mL) were seeded in T25 flasks. After 24 h, cells were transfected with an LP donor plasmid, a gRNA plasmid targeting the non-coding region (site T2), and a Cas9 plasmid at a ratio of 1:1:1 (w/w) using Lipofectamine 2000 (Invitrogen), according to the manufacturer’s instructions. To generate stable cell pools, 800 μg/mL hygromycin (Clontech, San Jose, CA) was used for the selection. The medium was changed every three days. After two weeks of selection, mCherry-positive/ZsGreen1-negative cell pools were sorted using FACS Aria II (BD Biosciences, San Jose, CA) with a 488 nm blue laser and 530/30 and 610/20 filters. Subsequently, stable cell lines were generated using limiting dilution method to a concentration of 0.3 cell/well into a 96-well plate. Clones were expanded and verified by 5’/3’-junction polymerase chain reaction (PCR), copy number, mRNA expression, and fluorescence level analyses (Supplementary Fig. S1).

### 2.5. Generation of a cell-based gRNA library

To ensure the coverage of 500 cells per gRNA, which can ensure sufficient representation of the gRNA library (Doench, 2018; Joung et al., 2017), the number of cells required for transfection was calculated based on the measured value of RMCE efficiency, which was 10.5% (Supplementary Fig. S2). One day before transfection, a total of 3.0 × 10^8^ cells were seeded at 0.5 × 10^6^ cells/mL in three 500 mL Erlenmeyer flasks (Corning) containing 200 mL of CD-CHO supplemented with 4 mM glutamine. On the day of transfection, cells were reseeded at 1.0 × 10^6^ cells/mL in twelve 125 mL Erlenmeyer flasks containing 50 mL of the medium. Cells were then transfected with the gRNA library and NLS-Bxb1 recombinase plasmids at a ratio of 3:1 (w/w) using a FreeStyle Max transfection reagent (Thermo Fisher), according to the manufacturer’s instructions. One day after transfection, an anti-clumping agent was added to the cells. Two days after transfection, cells were combined and sub-cultured into four 500 mL Erlenmeyer flasks containing 250 mL of the medium. Three days after transfection, cells were treated with 10 μg/mL puromycin (Sigma-Aldrich). Cell pools were passaged every three days with 10 μg/mL puromycin treatment. Fourteen days after transfection, recovered cell pools were subjected to Cas9 transfection, and a total of 5.6 × 10^7^ cells were used for genomic DNA extraction.

### 2.6. Generation of a knockout library cell pool

To ensure sufficient coverage, the number of cells required for transfection was calculated based on the measured Cas9 transfection efficiency, which was 20.4% (Supplementary Fig. S3). One day before transfection, a total of 3.0 × 10^8^ cells were seeded at 0.5 × 10^6^ cells/mL in three 500 mL Erlenmeyer flasks containing 200 mL of CD-CHO supplemented with 4 mM glutamine. On the day of transfection, cells were reseeded at 1.0 × 10^6^ cells/mL in twelve 125 mL Erlenmeyer flasks containing 50 mL of the medium. Cells were then transfected with a Cas9-BSD plasmid using the FreeStyle Max transfection reagent (Thermo Fisher), according to the manufacturer’s instructions. One day after transfection, cells were treated with 10 μg/mL blasticidin (Sigma-Aldrich). Two days after transfection, the medium was replaced with a fresh medium containing 10 μg/mL blasticidin. Four days after transfection, cells were recovered in a medium without blasticidin. Eight and sixteen days after transfection, a total of 5.6 × 10^7^ cells were prepared for genomic DNA extraction.

### 2.7. Hyperosmotic stress screening

To prepare a hyperosmolar medium (463 ± 4 mOsm/kg), 1.8 mL of 5 M NaCl (Sigma-Aldrich) was added to 118.2 mL of the standard medium (325 ± 1 mOsm/kg), which was the CD-CHO supplemented with 4 mM glutamine and 100X anti-clumping agent. Osmolality was measured using the Fiske Micro-Osmometer (Thermo Fisher Scientific). To sufficiently cover 500 cells per gRNA, a total of 6.0 × 10^7^ cells were seeded in triplicate at 0.5 × 10^6^ cells/mL in 500 mL Erlenmeyer flasks containing 120 mL of the standard or hyperosmolar medium and passaged every three days. After 21 days, a total of 5.6 × 10^7^ cells were used for genomic DNA extraction.

### 2.8. Preparation of next-generation sequencing (NGS) samples

Genomic DNA samples were extracted using the Exgene Blood SV kit (GeneAll Biotechnology, Seoul, South Korea), according to the manufacturer’s instructions. To prepare NGS samples, PCR was performed in a total volume of 50 μL with 4.0 μg genomic DNA per reaction using NEBNext Ultra II Q5 Master Mix (New England Biolabs, Ipswich, MA) (98 °C for 3 min; 22 cycles: 98 °C for 10 s, 60 °C for 30 s, 72 °C for 30 s; 72 °C for 5 min) using primers listed in Supplementary Table S2. PCR products were purified using a NucleoSpin Gel and PCR purification kit (Macherey-Nagel) and indexed using a TruSeq Nano DNA Library Prep kit (Illumina, San Diego, CA). The resulting library was quantified with a Qubit Flex Fluorometer (Thermo Fisher Scientific) using a dsDNA HS Assay kit (Thermo Fisher Scientific). Fragment size was determined using a 2100 Bioanalyzer Instrument (Agilent, Santa Clara, CA) and TapeStation D5000 (Agilent) and sequenced on a NextSeq 500 sequencer or a NextSeq 550 sequencer (Illumina).

### 2.9. NGS data analysis

For the analysis of gRNA fold-changes, raw FASTQ files were analyzed using Model-based Analysis of Genome-wide CRISPR/Cas9 Knockout (MAGeCK) (Li et al., 2014) and Platform-independent Analysis of PooLed screens using Python (PinAPL-Py) (http://pinapl-py.ucsd.edu/) (Spahn et al., 2017). MAGeCK v0.5.9.5 was run following the instructions (https://sourceforge.net/p/mageck/wiki/Home/#usage). The gene-test-fdr-threshold parameter was set to 0.01, and the other parameters were set to default. PinAPL-Py v2.9 was run using an adjusted robust rank aggregation (αRRA) ranking metric, an fdr_hb p-value adjustment method, and other default parameters. Top candidates for enriched gRNAs were ranked using the αRRA method and filtered using a p-value threshold of 0.01. A gene was considered to be “significant” if it was statistically significant at the gRNA level for at least half of the gRNAs from the total designed gRNAs targeting the gene.

### 2.10. Functional enrichment analysis

Gene ontology (GO) enrichment analysis was performed using the Database for Annotation, Visualization, and Integrated Discovery (DAVID) Knowledgebase v2022q2 (https://david.ncifcrf.gov/) (Huang da et al., 2009; Sherman et al., 2022). Pathway and process enrichment analyses were carried out using ontology resources from GO biological processes (Ashburner et al., 2000), Kyoto Encyclopedia of Genes and Genomes (Kanehisa and Goto, 2000), Reactome Gene Sets (Fabregat et al., 2018), and Molecular Signatures Database (Subramanian et al., 2005) using Metascape (https://metascape.org/) (Zhou et al., 2019). The network was visualized using Cytoscape v3.9.1 (Shannon et al., 2003).

### 2.11. Generation of knockout cell pools using all-in-one CRISPR/Cas9 plasmids

To generate a knockout cell pool in CHO-K1 cells, MCL was seeded at 1 × 10^6^ cells/mL in a 12-well plate containing 1 mL CD-CHO supplemented with 4 mM glutamine and transfected with all-in-one CRISPR/Cas9 plasmids targeting each gene using FreestyleMax, according to the manufacturer’s instructions. After 48 h, transfected cells were treated with 150 μg/mL blasticidin for three days and recovered for nine days without blasticidin. To generate knockout cell pools in CHO-mAb and CHO-bsAb cell lines, cells were seeded at 1 × 10^6^ cells/mL in a 6-well plate containing 3 mL SFM4Transfx-293 (HyClone) supplemented with 4 mM glutamine and transfected with all-in-one CRISPR/Cas9 plasmids targeting each gene using FreestyleMax according to the manufacturer’s instructions. After 48 h, transfected cells were treated with 75 μg/mL blasticidin for CHO-mAb and 150 μg/mL blasticidin for CHO-bsAb cell lines. After three days of blasticidin treatment, cells were recovered for nine days without blasticidin treatment.

### 2.12. Batch culture

CHO-K1 knockout cell pools were seeded at 0.5 × 10^6^ cells/mL in a 12-well plate containing 1 mL of the hyperosmolar medium used for screening and incubated at 37 °C, 85 % humidity, 5 % CO_2_, and 110 rpm. Cell viability and concentration were measured every two days. For CHO-mAb and CHO-bsAb knockout cell pools, hyperosmolar media (506.0 ± 3.6 and 463.7.0 ± 3.8 mOsm/kg, respectively) were prepared by adding 600 μL of 5M NaCl to 30 mL of culture media. CHO-mAb and CHO-bsAb knockout cell pools were seeded at 0.5 × 10^6^ cells/mL in 125 mL Erlenmeyer flasks with 30 mL of the hyperosmolar medium and incubated at 37 °C, 85 % humidity, 5 % CO_2_, and 110 rpm. Cell viability and density were measured daily. Culture supernatants were sampled daily and stored at −70 °C for further analysis.

### 2.13. Measurement of mAb and bsAb concentration

mAb and bsAb concentrations were measured using enzyme-linked immunosorbent assay, as previously described (Kim et al., 1998). The specific productivity was calculated from a plot of mAb and bsAb concentrations against the time integral values of viable cell concentration (VCC), as previously described (Renard et al., 1988).

### 2.14. Quantitative real-time polymerase chain reaction (qRT-PCR)

Total RNA was extracted using a Hybrid-R RNA extraction kit (GeneAll Biotechnology), and cDNA was synthesized using a high-capacity cDNA reverse transcription kit (Applied Biosystems, Foster City, CA), according to the manufacturer’s instructions. qRT-PCR was performed using iQ SYBR Green Supermix (Bio-Rad) on a CFX96 Real-Time System (Bio-Rad), as previously described (Noh et al., 2018). The relative expression levels were calculated using the ΔΔCT method and normalized to *Gapdh*. All primer sequences used in this study are listed in Supplementary Table S2.

### 2.15. Statistical analysis

Values are represented as the mean ± standard deviation. Data were analyzed using a two-tailed Student’s *t*-test, and the difference between the means was considered statistically significant at *P* < 0.05.

## 3. Results

### 3.1. Establishment of a virus-free, CHO-K1 genome-wide CRISPR knockout screening platform

A schematic illustration of the virus-free RMCE-based CRISPR knockout library platform is shown in Fig. 1. A CHO genome-wide CRISPR knockout library consisting of 111,651 gRNAs targeting 21,585 genes was designed, as described in the Materials and methods section (Supplementary Data File 1). gRNAs were synthesized and cloned into an RMCE donor plasmid, harboring a gRNA expression cassette and recombinase target sites. The cloned gRNA library plasmid was amplified and sequenced using NGS to determine the gRNA distribution. The coverage of the gRNA library plasmid was 99.9% and the skew ratio was 1.89, indicating sufficient representation with minimal bias. Next, to establish an RMCE platform in CHO-K1 cells, a landing pad harboring the mCherry gene and recombinase target sites was integrated into CHO-K1 cells. Based on homogeneous mCherry expression, clone #17 was selected, which will be referred to as “MCL” hereafter (Supplementary Fig. S1). To improve the RMCE efficiency of MCL, we generated an NLS-Bxb1 recombinase plasmid, which increased the efficiency by 1.6-fold compared to that of the control Bxb1 recombinase plasmid (Supplementary Fig. S4).

**Fig. 1.**
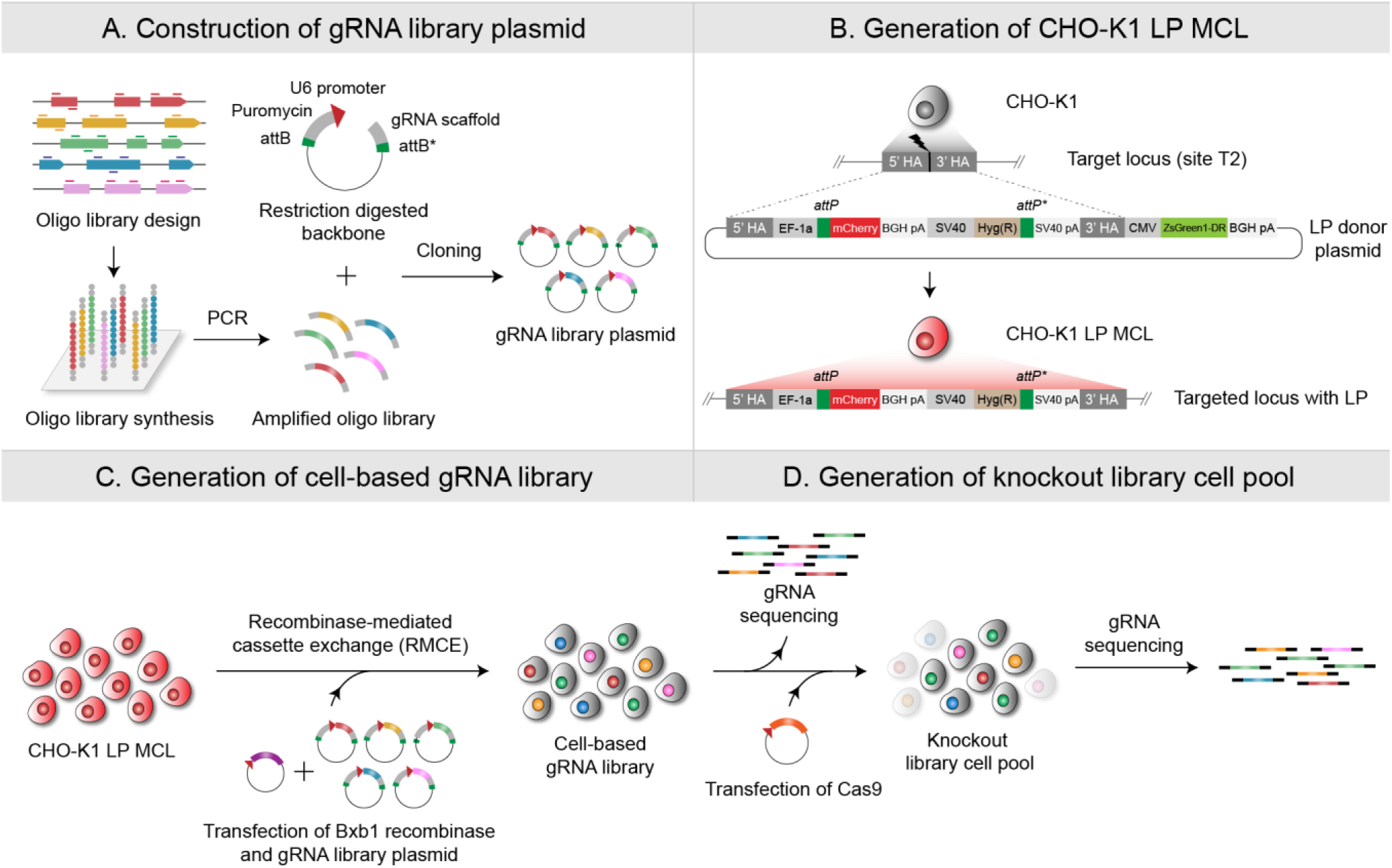
Establishment of a virus-free CHO-K1 genome-wide CRISPR knockout screening platform. (A) CHO-K1 CRISPR knockout guide RNAs (gRNAs) were designed and cloned into a gRNA scaffold backbone plasmid containing the puromycin resistance gene flanked by attB and attB^mut^ recombination sites. (B) CHO-K1 landing pad (LP) master cell line (MCL) harboring mCherry and hygromycin resistance genes flanked by attP and attP^mut^ recombination sites was generated via CRISPR/Cas9-based targeted integration. (C) CHO-K1 LP MCL was transfected with Bxb1 recombinase and gRNA library plasmid to generate a cell-based gRNA library. (D) CHO knockout library cell pool was generated via Cas9 plasmid transfection. To verify the representation of gRNA library in cell-based gDNA library and knockout library cells, gRNA sequences were sequenced using next-generation sequencing (NGS).

To introduce a gRNA library into MCL using RMCE, MCL was transfected with a gRNA library and NLS-Bxb1 recombinase plasmids, ensuring coverage of approximately 500 cells per gRNA. For enrichment of cells harboring the integrated gRNA library, the RMCE cell pool was treated with 10 μg/mL puromycin on day 3, and the puromycin selection was performed for 11 days (Fig. 2A and B). RMCE-positive cells were enriched from 16.4 to 99.4%, as evidenced by the percentage of mCherry-negative cells (Fig. 2C). To verify the representation of the gRNA library, gRNA sequences in the genomic DNA from the cell-based gRNA library were amplified and sequenced using NGS. The coverage of cell-based gRNA library was 99.7% and the skew ratio was 1.98, showing an even distribution similar to the plasmid library (Fig. 2D and E). Thus, the gRNA library was sufficiently represented in the cell-based gRNA library.

**Fig. 2.**
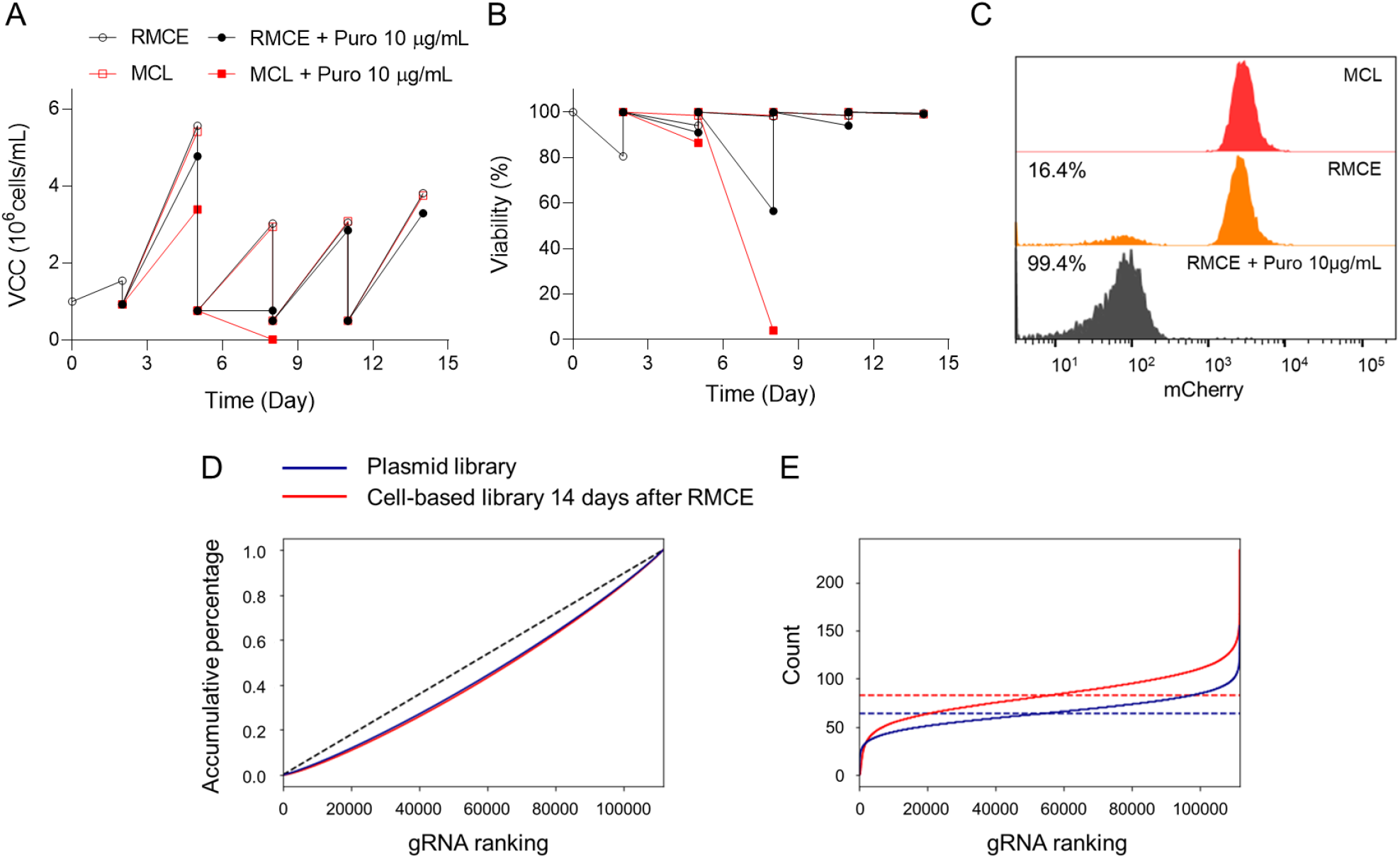
Generation of a cell-based gRNA library and representation of gRNA coverage. Profiles of (A) cell growth (viable cell concentration, VCC) and (B) viability of RMCE cell pool (black empty circle), RMCE cell pool with 10 μg/mL of puromycin (black full circle), MCL (red empty square), and MCL with 10 μg/mL of puromycin (red full square). On day 0, MCL was transfected with NLS-Bxb1 recombinase and gRNA library plasmids to obtain the RMCE cell pool. On day 2, the cells were sub-cultured. On day 3, RMCE cell pool and MCL control cells were treated with 10 μg/mL of puromycin for 11 days. (C) Flow cytometry analysis of the RMCE cell pool. Fourteen days after RMCE, cell populations expressing mCherry in MCL control (red), RMCE cell pool (orange), and RMCE cell pool with 10 μg/mL of puromycin (black) were measured. The percentage of mCherry-negative cells are shown. (D) Cumulative percentage of reads and (E) the number of read counts per gRNA in the plasmid library (blue solid line) and cell-based gRNA library 14 days after RMCE (red solid line). Dashed lines indicate the ideal models in the gRNA library.

Knockout library cells were generated by the transient transfection of Cas9-BSD plasmid, followed by three days of blasticidin selection to enrich the transfected cells (Fig. 3A). Eight days and sixteen days after Cas9 transfection, the recovered cells were harvested for NGS analysis. To assess the representation of the gRNA library, gRNA sequences in the genomic DNA from the knockout library cell pools were amplified and sequenced using NGS. As expected, Cas9 transfection perturbed the distribution of the gRNA library. The percentage of depleted gRNAs and genes gradually increased after Cas9 transfection, as shown in Fig. 3B and C, indicating that the genes essential for cell survival were affected. Sixteen days after Cas9 transfection, the generated knockout library cells were screened.

**Fig. 3.**
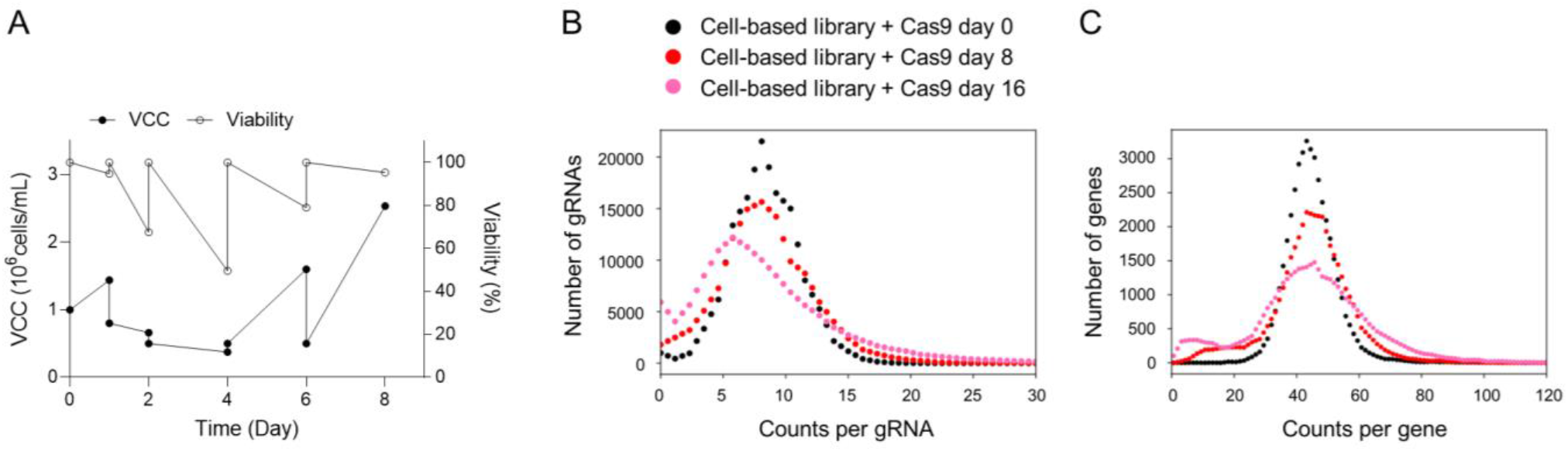
Generation of a knockout library cell pool and representation of library distribution. (A) Cell growth (viable cell concentration, VCC; full circle) and viability (empty circle) of the Cas9-transfected knockout library cell pool. On day 0, puromycin enriched RMCE cell pool was transfected with Cas9-BSD plasmid. On day 1, cells were treated with 10 μg/mL of blasticidin for three days. Read distributions of (B) gRNAs and (C) genes after Cas9 transfection in cell-based gRNA library. Cell based library (black), eight days after Cas9 transfection (red), and 16 days after Cas9 transfection (magenta).

### 3.2. Hyperosmotic stress screening

To determine the appropriate osmolality of the hyperosmolar medium for screening, knockout library cells were cultured in a hyperosmolar medium in which osmolality increased linearly by 50 mOsm/kg with NaCl addition (Supplementary Fig. S5A). Hyperosmolality negatively affected the cell growth and viability (Supplementary Fig. S5B and C). Osmolality of the hyperosmolar medium for screening was determined to be 460 mOsm/kg, which was the highest osmolality that the cells could tolerate (Supplementary Fig. S5D).

Knockout library cells were seeded at 0.5 × 10^6^ cells/mL in 1 L Erlenmeyer flasks containing 400 mL of the standard or hyperosmolar medium and passaged every three days (Fig. 4A). Cell cultures were performed in triplicate. Initially, cell growth was significantly suppressed in the hyperosmolar medium than in the standard medium (Fig. 4B). On day 9, the specific growth rate (*μ*) of knockout library cells in the standard medium was 0.71 ± 0.01 day^−1^, while that in the hyperosmolar medium was 0.14 ± 0.03 day^−1^ (Fig. 4C). However, the *μ* of cells in the hyperosmolar medium gradually increased and became saturated on day 21, while that in the standard medium remained constant during the culture. To evaluate the changes in gRNAs in standard and hyperosmolar media, cells were harvested on day 21 and subjected to NGS analysis.

**Fig. 4.**
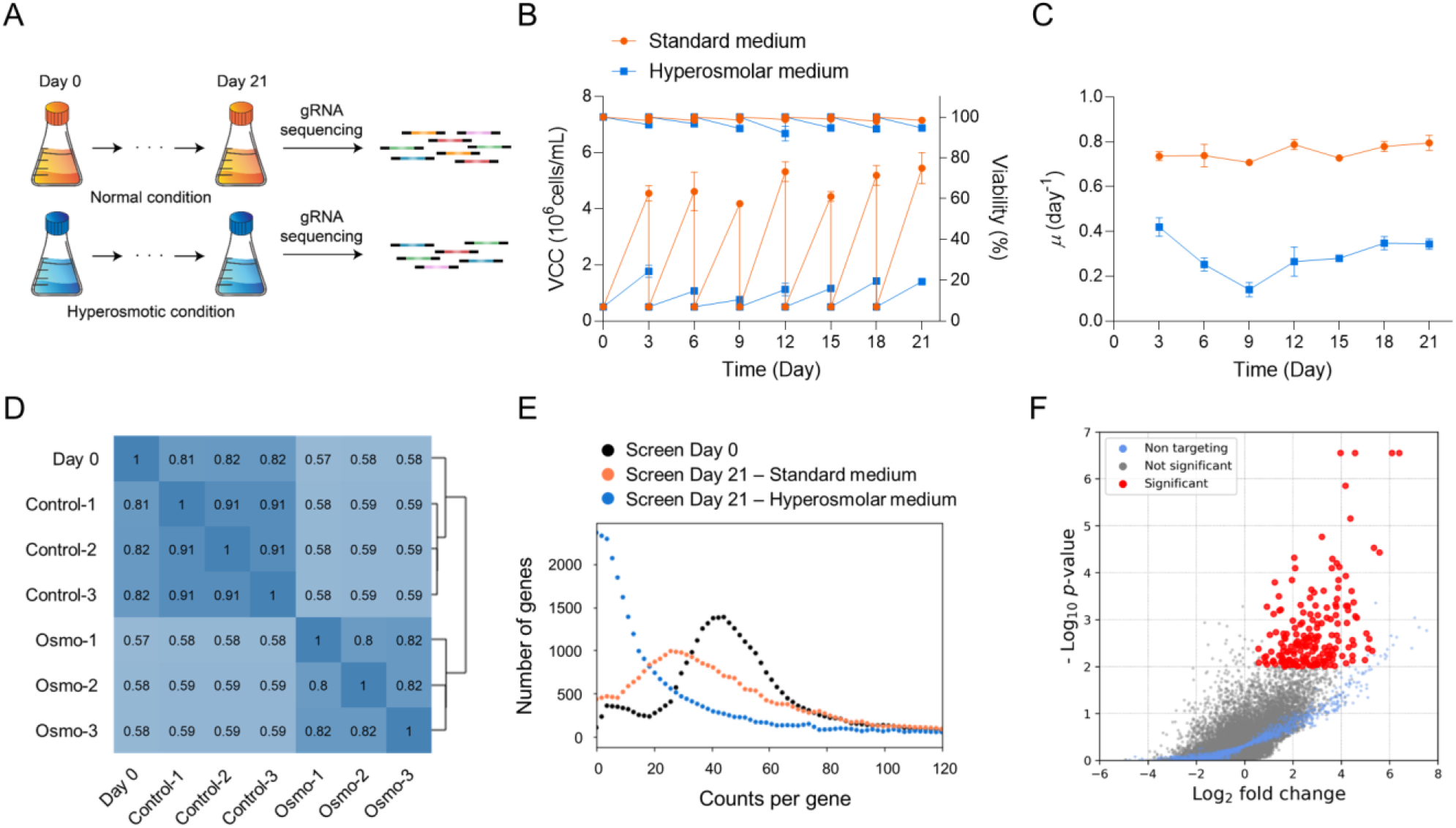
Hyperosmotic stress screening. (A) Schematic diagram illustrating hyperosmotic stress screening. Profiles of (B) cell growth (VCC), viability, and (C) *μ* in the standard medium (orange circle) and hyperosmolar medium (blue square). (D) Pearson correlation coefficient and sample hierarchical clustering of the gRNA read counts between samples of day 0 and triplicates of the standard medium (control) and hyperosmolar medium (osmo). (E) Read distribution of gRNA target genes. Day 0 (black), average of triplicates of standard medium (orange), and average of triplicates of hyperosmolar medium (blue). (F) Volcano plot showing the Log_2_ fold-change and minus Log_10_ *P*-value for each gene. Significantly enriched genes (*P*-value < 0.01) (red), non-targeting genes (blue), and not significant genes (gray).

To determine the gRNA abundance and distribution, computational analysis was conducted using MAGeCK (Supplementary Data File 2). The gRNA read counts of cells sampled on day 21 were compared to those of cells sampled on day 0. When correlated with cells sampled on day 0, cells sampled on day 21 in the hyperosmolar medium (correlation coefficient of 0.58) showed less correlation than cells sampled on day 21 in the standard medium (correlation coefficient of 0.82) (Fig. 4D). The first principal component obtained via principal component analysis showed clustering of cells on days 0 and 21 in the standard medium, while it showed separation with 60% of variation of cells on days 0 and 21 in the hyperosmolar medium (Supplementary Fig. S6). The variation can be explained by the perturbation in the read distribution, as the percentage of depleted gRNA target genes significantly increased after 21 days of cultivation in the hyperosmolar medium (Fig. 4E). Next, the fold-change in gRNAs between normal and hyperosmotic conditions was evaluated. The αRRA algorithm was used to rank genes by combining fold-change data from all gRNAs, and 180 significantly enriched genes were found on the screen (Fig. 4F). Taken together, hyperosmotic stress (460 mOsm/kg) provided sufficient selection pressure for the screen to generate perturbations in the cell pools.

### 3.3. Functional enrichment analysis

To characterize the enriched genes on the screen, GO enrichment analysis was performed using DAVID bioinformatics resources. GO enrichment analysis revealed the most significantly enriched GO terms for tRNA wobble uridine modification in biological processes and elongator holoenzyme complexes in cellular components (Fig. 5A and B). Both GO terms were composed of elongator acetyltransferase complex subunit (*Elp*) genes, which are required for tRNA modifications. In addition, biological processes of regulation of cell cycle, cellular components of the cytosol and mitochondria, and molecular functions of protein phosphatase-binding and methylated histone-binding were enriched (Fig. 5C).

**Fig. 5.**
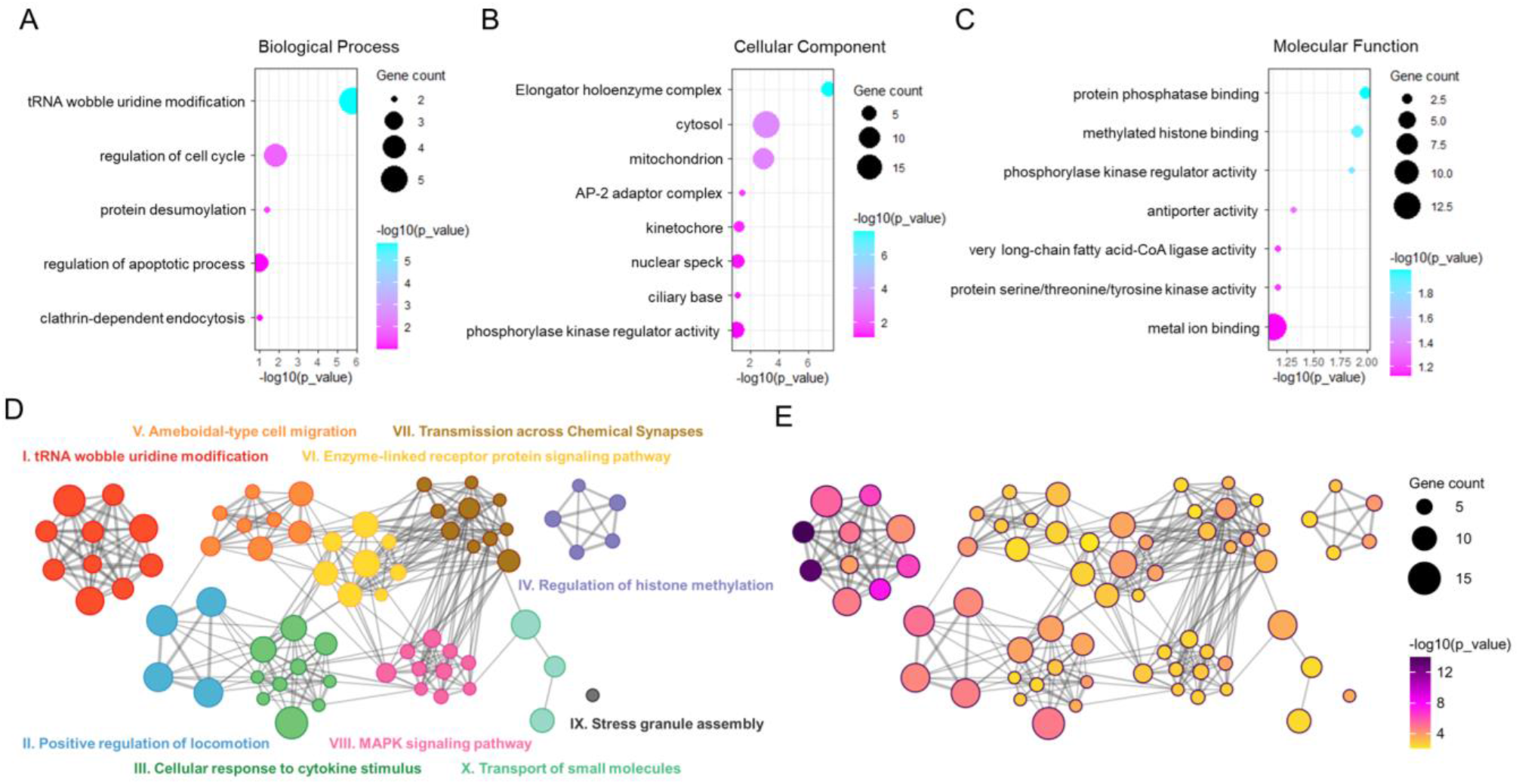
Functional enrichment analysis using enriched genes to identify hyperosmotic stress-responsive gene networks. Bubble plots showing enriched GO terms in (A) biological processes, (B) cellular components, and (C) molecular functions. The size of each circle indicates the number of genes that are enriched in the GO term. Network plots showing pathway and process enrichment analysis colored by (D) cluster and (E) *P*-value. Each node represents an individual enriched term. A connection between nodes represents Kappa similarity above 0.3, forming a network cluster. The node size indicates the number of genes enriched in the node.

To identify hyperosmotic stress-responsive pathways on a genome-wide scale, pathway and process enrichment analyses were conducted using Metascape (Zhou et al., 2019). 10 functional clusters of enriched biological terms were identified, and the relationships among the terms were visualized as a network plot (Fig. 5D). A cluster related to tRNA wobble uridine modifications was significantly enriched (Fig. 5E). Clusters related to the regulation of cell migration, cellular response to cytokine stimulus, regulation of histone methylation, mitogen-activated protein kinase (MAPK) signaling pathways, and transportation were also enriched. Raw data for pathway and process enrichment analyses are summarized in Supplementary Data File 3.

### 3.4. Validation of candidate genes in CHO-K1 cells

To generate a robust data-set and prioritize genes from 180 significantly enriched genes derived from MAGeCK, computational analysis was conducted using another analysis tool, PinAPL-Py, and 58 significantly enriched genes were identified (Supplementary Data File 4). Among the 58 genes, 52 genes were enriched in both MAGeCK and PinAPL-Py analyses. To prioritize genes, the top 100 ranking genes were filtered using the MAGeCK aRRA score, and 32 genes were sorted (Fig. 6A). *Fastkd1* and *Zfr* were the highest scoring genes, and all gRNAs targeting *Fastkd1* and *Zfr* showed significant enrichment in the screen. GO enrichment analysis was conducted to characterize the functions of these 32 genes. Three enriched GO terms of biological processes were found to be tRNA wobble uridine modification, regulation of mRNA stability, and small molecule catabolic processes (Fig. 6B). In addition to the three enriched GO terms, various gene functions, such as ion transport, regulation of the cell cycle, endocytosis, and cell junctions were identified. Detailed functions of the 32 genes are summarized in Table 1.

**Fig. 6.**
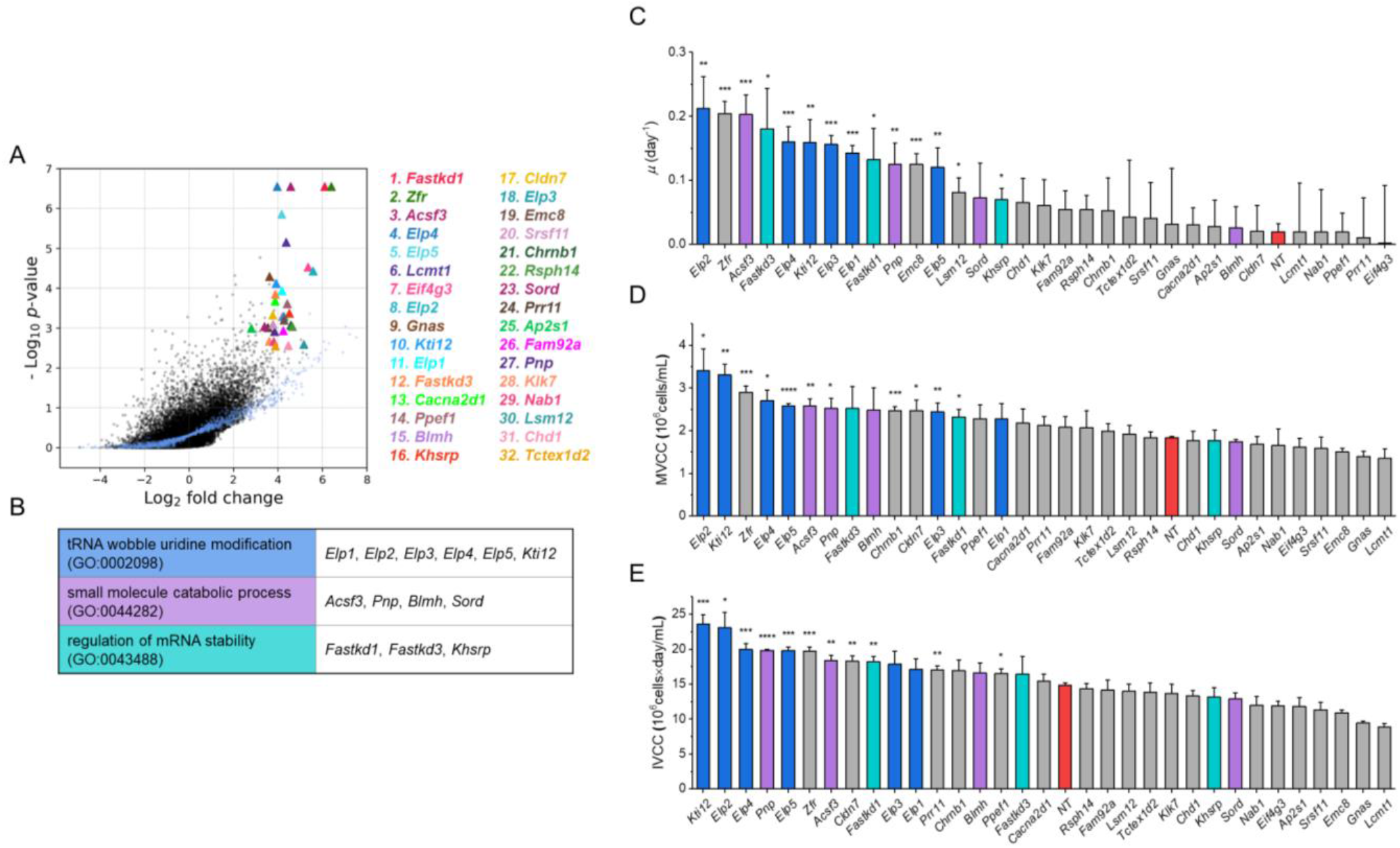
Validation of the candidate genes. (A) Log_2_ fold-change and minus Log_10_ *P*-value in MAGeCK analysis for the top-scoring 32 candidate genes. The number in front of the gene name indicates its ranking in the analysis. Genes are indicated by triangles with labeled color. Non-targeting genes are indicated by blue dots and not significant genes are indicated by black dots. (B) Enriched GO terms and genes. (C) *μ*, (D) MVCC, and (E) IVCC of the 32 candidate gene knockout pools. *μ* was calculated based on the values of VCC during the exponential phase (days 2-6). Columns in blue represent genes in the GO term of tRNA wobble uridine modification, in purple represent genes in the GO term of small molecule catabolic process, in cyan represent genes in the GO term of regulation of mRNA stability, and in red represent the NT control. Asterisks (*) indicate the significant difference compared to the NT control. Error bars in the plot represent the standard deviations of three biological replicates. An unpaired two-tailed *t*-test was used to determine the significance of the mean difference. \**P* < 0.05, \*\**P* < 0.01, \*\*\**P* < 0.001, and \*\*\*\**P* < 0.0001.

**Table 1.**
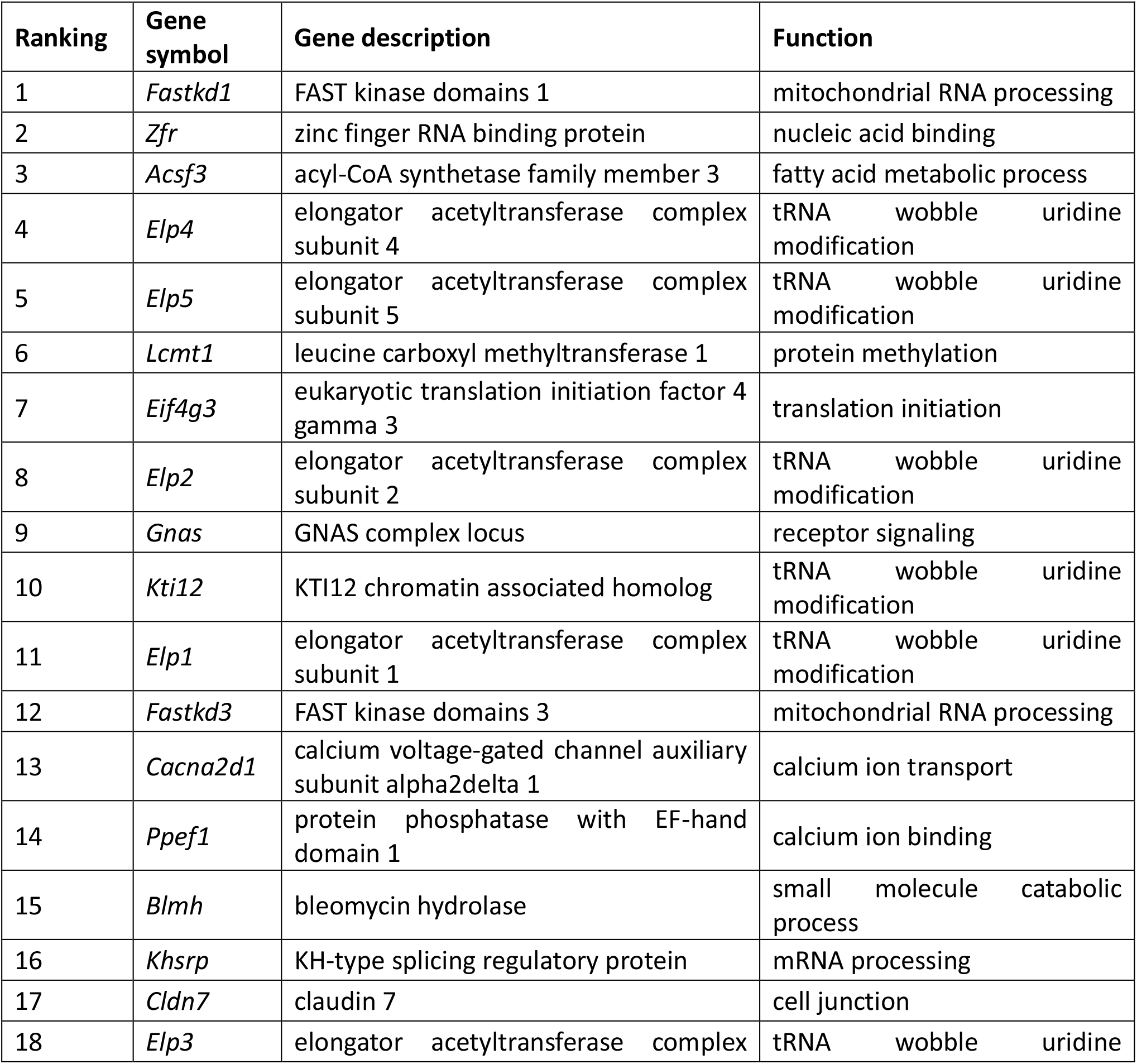
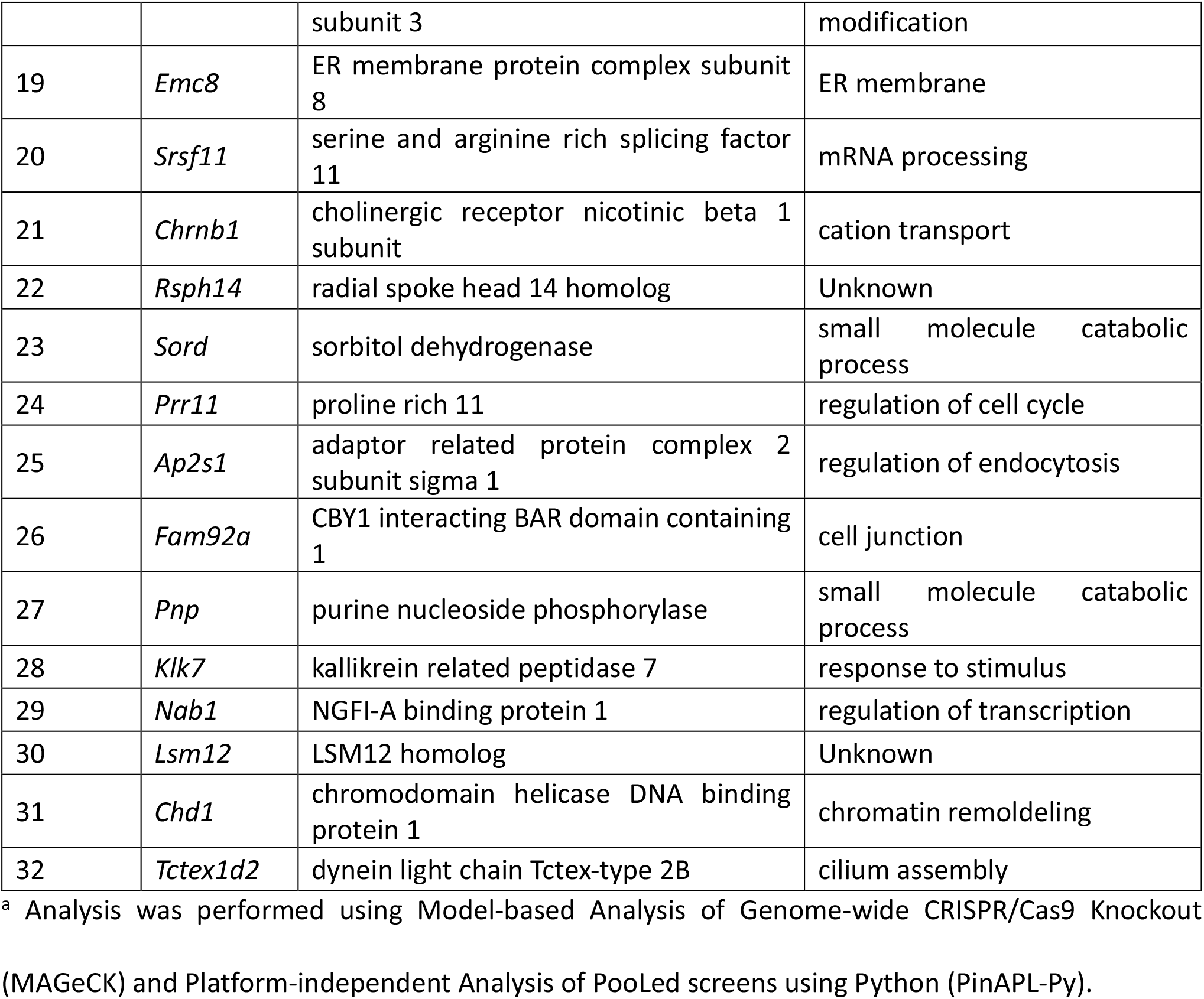
List of 32 candidate genes significantly enriched in the analysis.^a^

To verify the screening results, knockout cell pools for the 32 candidate genes were generated using all-in-one CRISPR/Cas9 plasmids. The knockout efficiency of the all-in-one CRISPR/Cas9 system was estimated by measuring the percentage of mCherry-negative cells in MCL using an mCherry-targeting all-in-one CRISPR/Cas9 plasmid and increased to 97.0% via blasticidin selection (Supplementary Fig. S7). A total of 32 knockout cell pools were cultured for 10 days in a 12-well plate containing the hyperosmolar medium used for screening. VCCs were measured every other day. Cell cultures were performed in three separate times.

Overall, 29 out of the 32 candidate gene knockout cell pools showed increased *μ* or maximum VCC (MVCC) in the hyperosmolar medium (Fig. 6C and D). In addition, eight of the 29 candidate gene knockout cell pools showed a significant increase in both *μ* and MVCC. (Fig. 6C and D). Interestingly, five (*Fastkd1*, *Zfr*, *Acsf3*, *Elp4*, *Elp5*) of the eight candidate genes were the most highly enriched genes in the screen.

*Kti12* knockout pool showed the highest time integral of VCC (IVCC) of 23.6 ± 1.4 × 10^6^ cells/mL·day, which is 1.6-fold higher than that of the NT control pool (Fig. 6E). In addition, the majority of candidate gene knockout pools showed higher viability than the NT control pool on day 10 (Supplementary Fig. S8). *Elp4* knockout pool showed the highest viability of 80% on day 10, whereas the viability of the NT control pool was 42% on day 10. Thus, knockout validation narrowed the candidate genes from the 32 candidate genes screened using hyperosmotic stress screening. Seven genes, whose knockout showed the highest increase in both MVCC and IVCC, were further validated. (MAGeCK) and Platform-independent Analysis of PooLed screens using Python (PinAPL-Py).

### 3.5. Assessment of gene knockout in mAb-producing rCHO cell lines

To further investigate the effects of target gene knockout in mAb-producing rCHO cell lines, knockout cell pools for seven candidate genes (*Zfr*, *Acsf3*, *Elp2*, *Elp4*, *Elp5*, *Kti12*, and *Pnp*) were generated using all-in-one CRISPR/Cas9 plasmids. Knockout of each target gene was verified via mRNA expression analysis, which showed decreased mRNA expression levels for each target gene (Supplementary Fig. S9). NT control and seven candidate gene knockout pools were cultured in the hyperosmolar medium, and VCCs and mAb concentrations were measured on day 6. Among the seven candidate gene knockout pools, *Zfr* and *Pnp* knockout pools showed significant increase in both cell growth and mAb productivity (Supplementary Fig. S10). To assess the growth profiles of *Zfr* and *Pnp* knockout pools in the hyperosmolar medium, NT control and *Zfr* and *Pnp* knockout cell pools were cultured in 125 mL Erlenmeyer flasks with 30 mL of hyperosmolar medium (506.0 ± 3.6 mOsm/kg). VCC was measured daily. Experiments were performed in three separate times.

Compared to the NT control, *Zfr* and *Pnp* knockout cell pools showed a higher VCC during culture (Fig. 7A). The viability of NT control cell pools dropped to 77.5 ± 3.5 % on day 2, while that of *Zfr* and *Pnp* knockout pools remained above 95% (Fig. 7B). IVCCs of *Zfr* knockout (27.8 ± 1.8 × 10^6^ cells/mL·day) and *Pnp* knockout (25.4 ± 0.9 × 10^6^ cells/mL·day) cells were 1.5 and 1.4-fold higher than that of NT (18.0 ± 1.1 × 10^6^ cells/mL·day) cells, which resulted in increased mAb production (Fig. 7C and D). The maximum mAb concentrations of *Pnp* knockout (429.4 ± 25.8 mg/L) and *Zfr* knockout (384.7 ± 13.8 mg/L) cells were 1.6- and 1.5-fold higher than that of NT (265.1 ± 15.8 mg/L) cells (Fig. 7C). However, the specific productivities (*q*_mAb_) of the knockout cell pools were not significantly different from that of the NT control cells (Fig. 7E). *Zfr* and *Pnp* knockout pools were also cultured in the standard medium, but no significant differences were observed compared to the NT control cells (data not shown). Therefore, knockout of *Zfr* and *Pnp* conferred osmotic stress resistance under hyperosmotic conditions and increased the cell growth and mAb production in CHO cells.

**Fig. 7.**
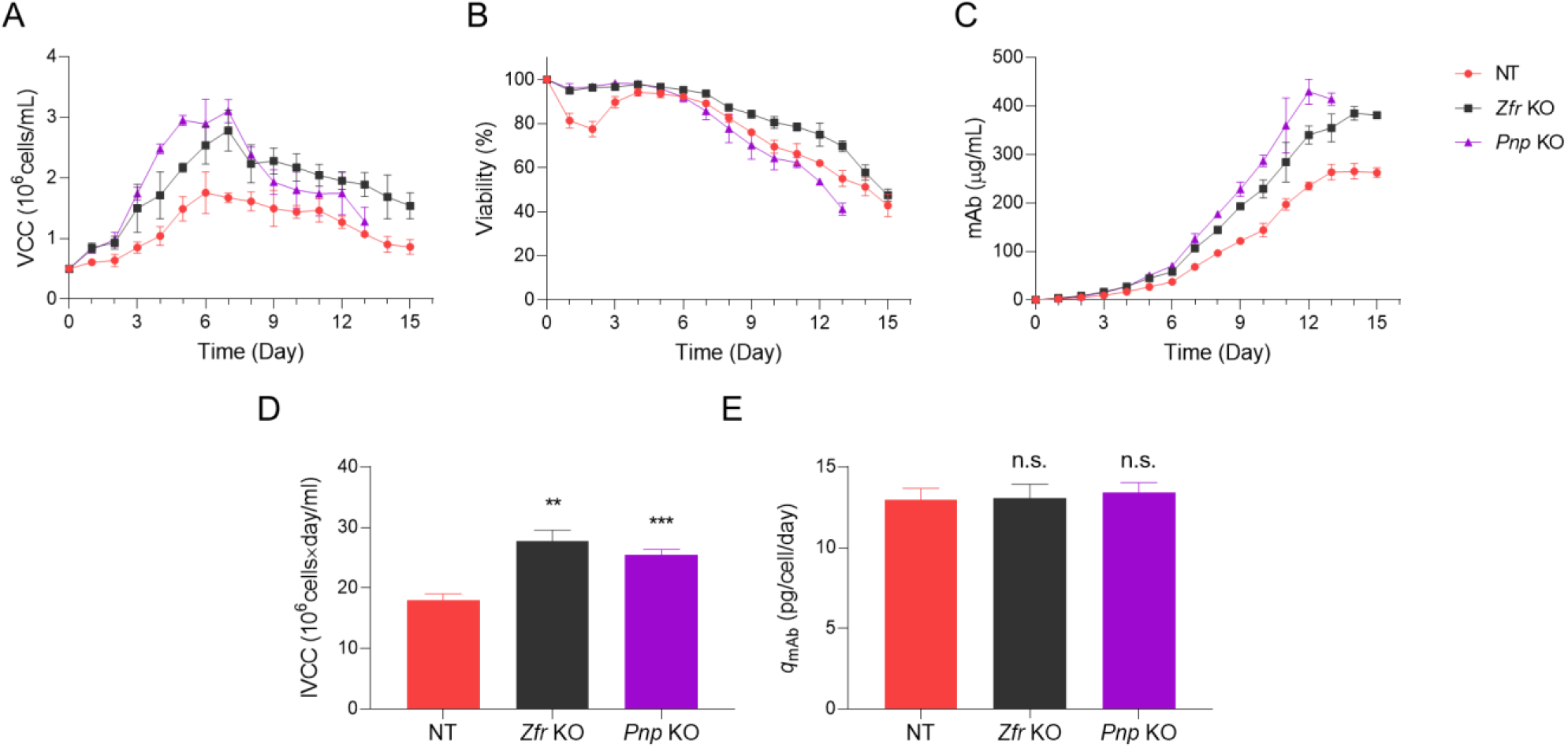
Batch cultures of *Zfr* and *Pnp* knockout CHO-mAb pools. Profiles of (A) cell growth (VCC), (B) viability, (C) mAb concentration, (D) IVCC, and (E) *q*_mAb_ of *Zfr* and *Pnp* knockout CHO-mAb pools and NT control pool in the hyperosmolar medium. Asterisks (*) indicate the significant difference compared to the NT control. Error bars in the plot represent the standard deviations of three biological replicates. An unpaired two-tailed *t*-test was used to determine the significance of the mean difference. n.s. *P* > 0.05, \*\**P* < 0.01, and \*\*\**P* < 0.001.

### 3.6. Assessment of gene knockout in bsAb-producing CHO-S cell lines

To determine whether knockout of *Zfr* and *Pnp* confer osmotic stress resistance in rCHO cell lines producing different antibody products generated from different lineages, CHO-S cell lines producing bsAbs were transfected with the NT control and *Zfr* and *Pnp* targeting all-in-one CRISPR/Cas9 plasmids. The relative mRNA expression levels of *Zfr* and *Pnp* in *Zfr* and *Pnp* knockout cell pools, respectively, showed successful gene knockout of each target gene (Supplementary Fig. S11). Knockout cell pools were cultured in 125 mL flasks with 30 mL of hyperosmolar medium (463.7 ± 3.8 mOsm/kg). VCC was measured daily. Experiments were performed in three separate times.

*Zfr* and *Pnp* knockout cell pools showed a higher VCC than the NT control cells during culture (Fig. 8A). IVCCs of *Zfr* knockout (24.5 ± 0.9 × 10^6^ cells/mL·day) and *Pnp* knockout (23.8 ± 0.8 × 10^6^ cells/mL·day) cells were 1.2-fold higher than that of NT (20.1 ± 0.6 × 10^6^ cells/mL·day) cells (Fig. 8D). The maximum bsAb concentrations of *Zfr* knockout (134.3 ± 1.6 mg/L) and *Pnp* knockout (123.4 ± 2.6 mg/L) cells were 1.3- and 1.2-fold higher than that of NT (100.4 ± 3.6 mg/L) cells (Fig. 8C). However, the *q*_mAb_ of knockout cell pools was not significantly different from that of the NT control cells (Fig. 8E). Therefore, knockout of *Zfr* and *Pnp* conferred osmotic stress resistance and increased the cell growth and bsAb production in CHO cells.

**Fig. 8.**
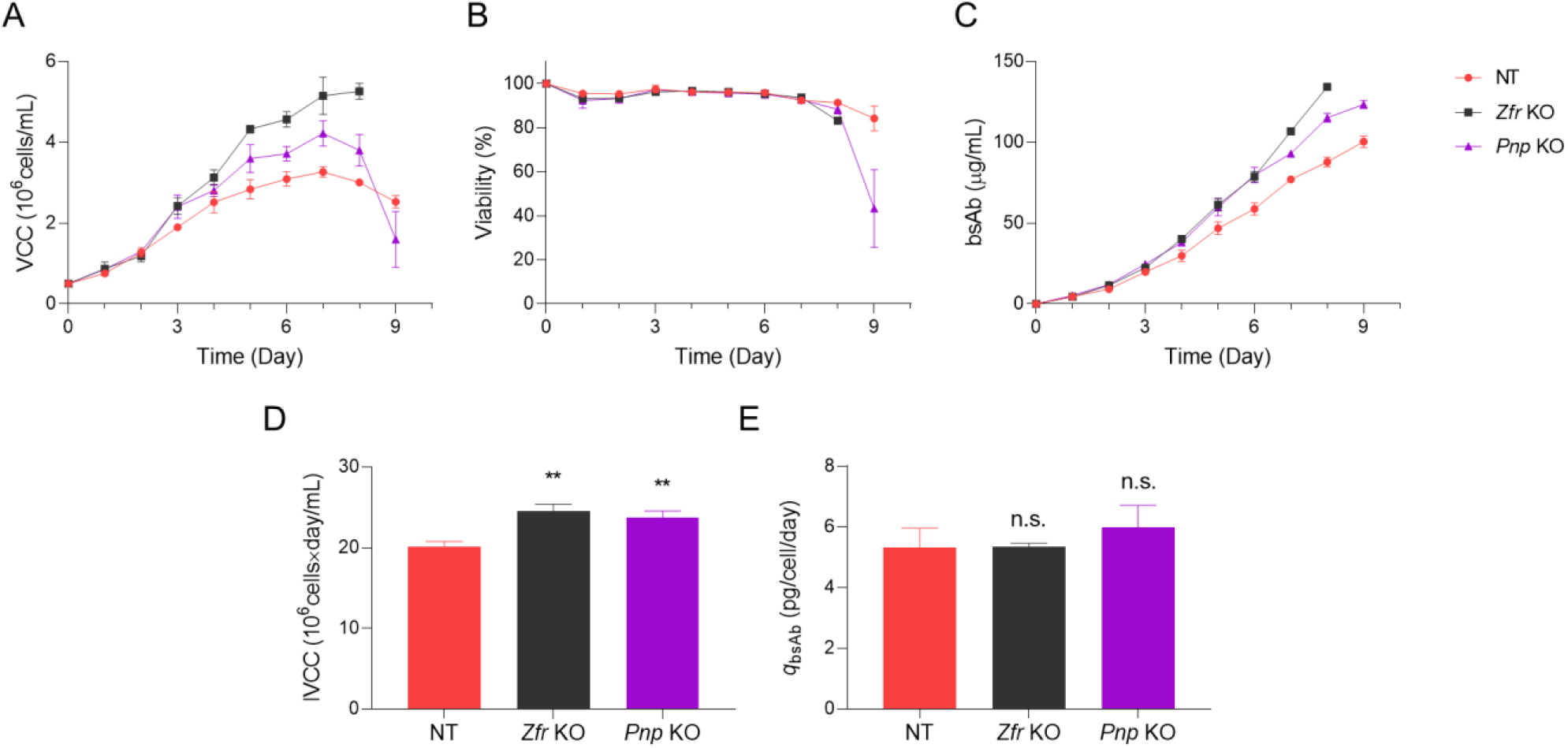
Batch cultures of *Zfr* and *Pnp* knockout CHO-bsAb pools. Profiles of (A) cell growth (VCC), (B) viability, (C) mAb concentration, (D) IVCC, and (E) *q*_mAb_ of *Zfr* and *Pnp* knockout CHO-bsAb pools and NT control pool in the hyperosmolar medium. Asterisks (*) indicate the significant difference compared to the NT control. Error bars in the plot represent the standard deviations of three biological replicates. An unpaired two-tailed *t*-test was used to determine the significance of the mean difference. n.s. *P* > 0.05, and \*\**P* < 0.01.

## 4. Discussion

Being powerful mammalian expression platforms for the production of therapeutic proteins, including mAbs, CHO cells have been engineered to achieve high productivity and quality of therapeutic proteins through knowledge-based approaches. However, empirical considerations for finding novel targets for desired attributes have marginal effects, necessitating the development of new strategies. Advances in CRISPR/Cas9 technology have generated a wealth of tools for genome-scale approaches, such as genetic screening, which enables unbiased dissection of genes with relevant phenotypes. Here, genome-wide CRISPR/Cas9 screening was performed in CHO cells to identify novel targets associated with industrially relevant hyperosmotic stress conditions.

Most of the current experimental designs in CRISPR/Cas9 screening involve lentiviral transduction of the gRNA library into Cas9 expressing cells, however, this can result in spurious gene editing of Cas9 guided by multiple gRNAs (Xiong et al., 2021). In addition, Cas9 expression can result in nuclease-induced cellular toxicity due to the DNA damage response (Morgens et al., 2017; Tycko et al., 2019). To circumvent these issues, RMCE-based gRNA integration followed by a transient Cas9 expression platform was previously established, demonstrating high efficiency both in single integration of gRNA and in Cas9 editing (Xiong et al., 2021). Using this platform, even distribution of gRNAs with high coverage was accomplished in our cell-based library (Fig. 2D), and knockout library cells were subsequently generated (Fig. 3).

Pooled screening requires a suitable selection pressure that can lead to perturbation of cell proliferation or viability, such that enriched or depleted mutants can be effectively discriminated. Osmotic tolerance differs among cell lines (Ryu et al., 2001). Thus, the appropriate hyperosmolar medium for each rCHO cell line was determined throughout the osmolality test (Supplementary Fig. S5; Fig. S12). Consequently, the hyperosmolar medium used for screening sufficiently generated perturbations in the cell pools and elicited significantly enriched and depleted genes (Fig. 4).

Hyperosmotic stress inhibits cell proliferation and induces apoptosis by activating signaling pathways mediated by MAPKs, including p38 and c-Jun N-terminal kinase (Zhou et al., 2016). As expected, functional enrichment analysis of genes enriched in the screen identified enriched terms of cell cycle arrest and the MAPK signaling pathway (Fig. 5). Representatively, *Pten*, *Nf2*, *Mapk14*, and *Traf3* have been identified. These genes are known to inhibit cell proliferation, and their knockout promotes cell proliferation (Brandmaier et al., 2017; Gurusamy et al., 2020; Xiao et al., 2005; Zhou et al., 2021). Additionally, hyperosmotic stress induces cell volumetric changes that accompany various alterations in transporters across the membrane (Hoffmann et al., 2009). We also found enriched terms of transporters, which included genes, such as chloride voltage-gated channel (*Clcn4*), calcium-activated cation channel (*Trpm4*), and solute carrier gene family. In particular, *Trpm4* can generate osmotic gradients, and knockdown of *Trpm4* gene attenuates cell swelling, preventing negative consequences in astrocytes (Stokum et al., 2018). Interestingly, while most functional terms were connected to form a network, the term tRNA wobble modification was separated but showed the most significant enrichment in functional analyses. Upon stress conditions, cells have been shown to support efficient translation of stress-responsive genes favoring rare tRNAs with wobble-pairing by altering tRNA abundance and modification (Frenkel-Morgenstern et al., 2012; Torrent et al., 2018), and the alteration of tRNA modifications contributes to a translational regulatory mechanism in stress response (Huang and Hopper, 2016).

Computational analyses identified 32 top-scoring genes, and the majority of knockout cell pools for each candidate gene showed hyperosmotic stress-resistance in CHO-K1 cells, validating the screening result (Fig. 6). Notably, knockout of *Fastkd1* and *Zfr*, which were the highest-scoring genes with significant enrichment of all target gRNAs, significantly increased the cell growth under hyperosmotic stress in CHO-K1 cells. In addition, perturbations of genes related to tRNA wobble modification (*Elp1*, *Elp2*, *Elp3*, *Elp4*, *Elp5*, and *Kti12*), which showed significant enrichment in functional analysis, resulted in high *μ* and MVCCs in CHO-K1 cells. While the hyperosmotic stress-resistant effect of knockout of *Zfr* and *Pnp* persisted in CHO-mAb cell lines, consistent phenotypes were not reproduced for some genes related to tRNA wobble modification (Supplementary Fig. S10). This can be explained by the dynamics of tRNA pools and variations of stress-responsive genes in various cell types, thereby causing to variations in cell lines (Gingold et al., 2012).

Hyperosmolality can increase *q*_p_ in CHO cells to different extents in different cell lines (Ryu et al., 2000); therefore, we investigated the effects of gene perturbations upon hyperosmotic stress in CHO cells of different origins, producing different products. Notably, perturbations of *Zfr* and *Pnp* genes in CHO-mAb and CHO-bsAb cell lines also increased the cell growth under hyperosmotic culture conditions and enhanced the product titers. However, this improvement was not attained by a substantial increase in *q*_p_ (Fig. 7E; Fig. 8E). Because hyperosmotic stress leads to an increase in *q*_p_ concomitant with cell cycle arrest, the enhancement of *q*_p_ may also be diminished by mitigating growth inhibition. PNP is an enzyme that converts inosine to hypoxanthine in the purine degradation pathway, and inosine enhances the cell proliferation of highly proliferating cells (Soares et al., 2015; Yin et al., 2018). Therefore, we speculate that blockage of inosine conversion by the knockout of *Pnp* enhanced the cell proliferation in this study. ZFR is a DNA- and RNA-binding protein; however, its biological functions remain unknown. Hence, further investigations of the roles of ZFR should be performed in future studies.

In conclusion, genome-wide CRISPR/Cas9 screening was performed in CHO-K1 cells, and novel genes and functional clusters associated with hyperosmotic stress were identified. Knockout of target genes increased the cell growth under hyperosmotic culture conditions, thereby enhancing the productivity. Our findings demonstrate the beneficial values of the screening platform in providing novel insights on hyperosmotic stress and identification of novel targets for rational cell engineering on a genome-wide scale.

## Abbreviations

bsAb: bispecific antibody
Cas9: CRISPR-associated protein 9
CHO: Chinese hamster ovary
CRISPR: clustered regularly interspaced short palindromic repeats
FACS: fluorescence-activated cell sorting
GO: Gene ontology
gRNA: guide RNA
LP: landing pad
mAb: monoclonal antibody
MCL: master cell line
NGS: next-generation sequencing
PCR: polymerase chain reaction
qRT-PCR: quantitative real-time polymerase chain reaction
RMCE: recombinase-mediated cassette exchange

## Author statement

**Su Hyun Kim**: Conceptualization, Methodology, Validation, Investigation, Writing – original draft, Writing – review & editing; **Seunghyeon Shin**: Validation, Investigation, Writing – review & editing; **Minhye Baek**: Validation, Investigation, Writing – review & editing; **Kai Xiong**: Methodology, Software, Validation, Resources; **Karen Julie la Cour Karottki**: Software, Formal analysis, Resources; **Hooman Hefzi**: Software, Formal analysis, Resources; **Lise Marie Grav**: Resources; **Lasse Ebdrup Pedersen**: Software, Resources; **Helene Faustrup Kildegaard**: Conceptualization; **Nathan E. Lewis**: Conceptualization, Formal analysis, Resources; **Jae Seong Lee**: Conceptualization, Methodology, Validation, Project administration, Funding acquisition, Writing – review & editing; **Gyun Min Lee**: Conceptualization, Methodology, Validation, Project administration, Funding acquisition, Supervision, Writing – review & editing

## Acknowledgments

The authors would like to thank Yeonwoo Kim for assistance with cell line maintenance and data processing in this study. Flow cytometry and NGS were performed at the Bio Core facilities of the Korea Advanced Institute of Science and Technology (KAIST). This research was supported by the Samsung Research Funding Center of Samsung Electronics under Project number SRFC-MA1901-09.

## Appendix A. Supplementary data

Supplementary data to this article can be found online at

