## Supplementary data for "Identification of hyperosmotic stress-responsive genes in Chinese hamster ovary cells via genome-wide virus-free CRISPR/Cas9 screening"

---

---

* Corresponding author. KAIST, Republic of Korea.

** Corresponding author. Ajou University, Republic of Korea

**Supplementary Figures**

Supplementary Figure S1. Generation of CHO-K1 landing pad (LP) master cell lines (MCLs) via CRISPR/Cas9-mediated integration.

Supplementary Figure S2. RMCE efficiency in an Erlenmeyer flask.

Supplementary Figure S3. Cas9 transfection efficiency.

Supplementary Figure S4. Determination of RMCE efficiency using Bxb1 recombinase and NLS-Bxb1 recombinase plasmids.

Supplementary Figure S5. Effects of osmolality on cell growth.

Supplementary Figure S6. Principal component analysis (PCA).

Supplementary Figure S7. Measurement of mCherry knockout efficiency in CHO-K1 MCL using an all-in-one CRISPR/Cas9 plasmid.

Supplementary Figure S8. Viabilities of the 32 candidate gene knockout cell pools.

Supplementary Figure S9. Relative mRNA expression levels of each target gene in CHO-mAb knockout cell pools.

Supplementary Figure S10. Assessment of gene knockout in CHO-mAb cell line.

Supplementary Figure S11. Relative mRNA expression levels of each target gene in CHO-bsAb knockout cell pools.

Supplementary Figure S12. Effects of osmolality on CHO-mAb and CHO-bsAb cell lines.

**Supplementary Tables**

Supplementary Table S1. Plasmid information.

Supplementary Table S2. Primer information.

Supplementary Table S3. gRNA information.

**Supplementary Data Files**

Supplementary Data File 1. CHO-K1 whole genome knockout gRNA library.

Supplementary Data File 2. Result of computational analysis using MAGeCK.

Supplementary Data File 3. Result of pathway and process enrichment analysis using Metascape.

Supplementary Data File 4. Result of computational analysis using PinAPL-Py.

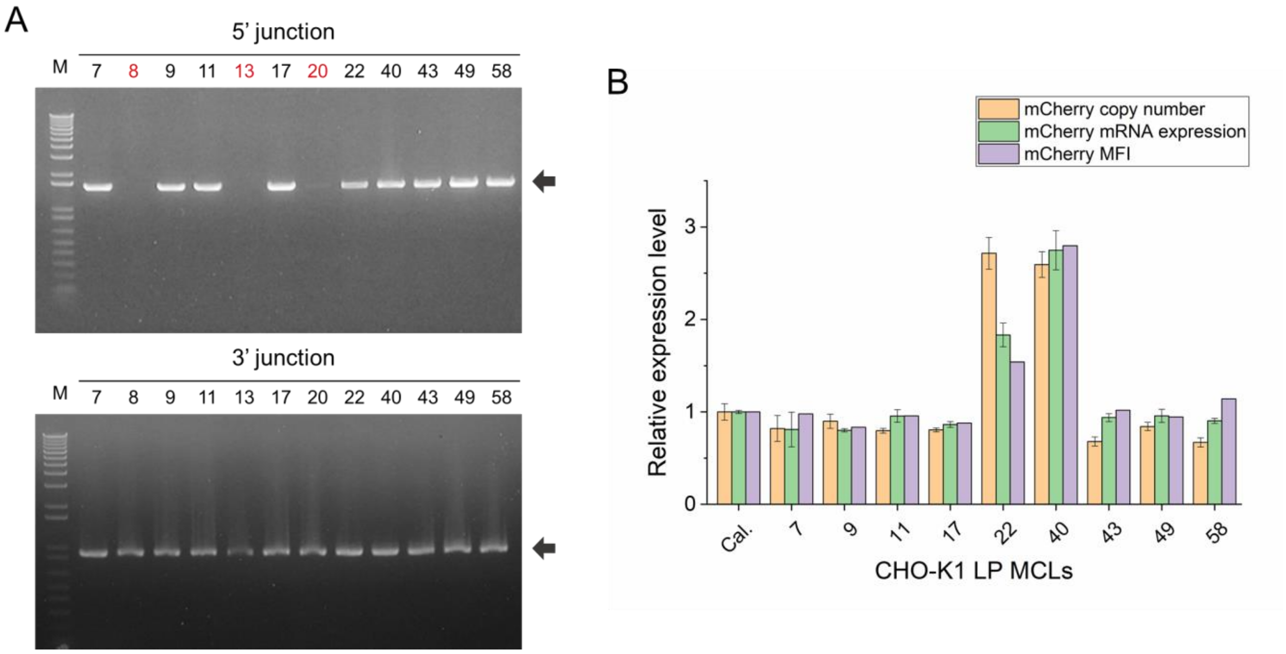


**Supplementary Fig. S1.** Generation of CHO-K1 landing pad (LP) master cell lines (MCLs) via CRISPR/Cas9-mediated integration. (A) Verification of targeted integration at the site T2 locus in 12 clones via 5'/3'-junction polymerase chain reaction (PCR). Negative clones are shown in red, and black arrows indicate the expected size of the PCR amplicons. (B) Measurement of the relative copy number, mRNA expression, and mean fluorescence intensity (MFI) of mCherry gene. CHO-K1 calibrator cell line (Cal.), which contains one mCherry gene in the site A locus, was used as a reference. Copy number, mRNA expression, and flow cytometry analyses were conducted, as previously described (Grav et al., 2018).

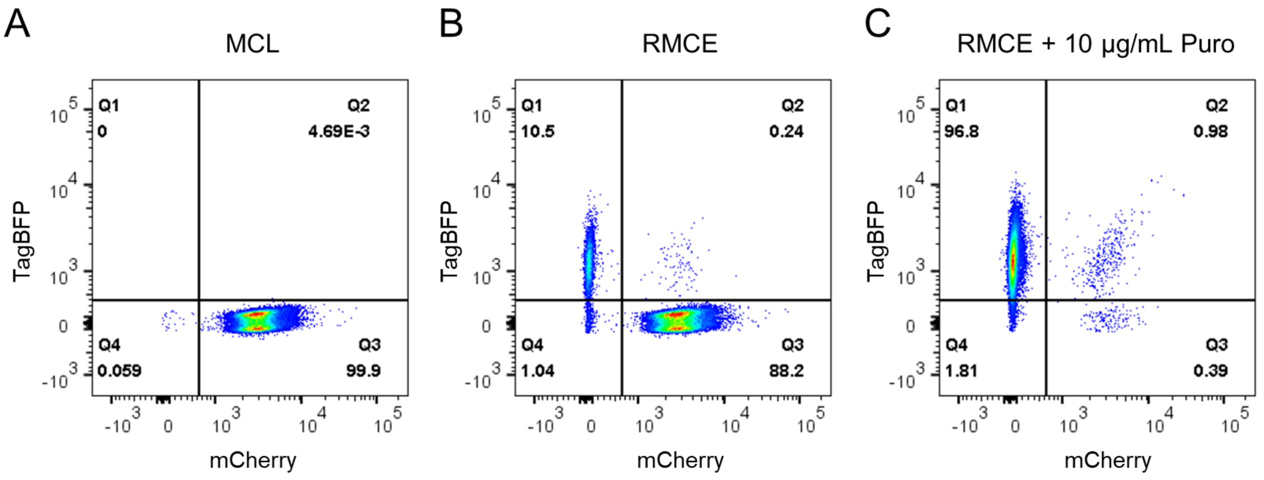


**Supplementary Fig. S2.** RMCE efficiency in an Erlenmeyer flask. Flow cytometry analysis of mCherry and TagBFP expression in (A) MCL, (B) 11 days after RMCE, and (C) 11 days after RMCE with 10 μg/mL of puromycin. MCL was seeded at a concentration of $1.0 \times 10^6$ cells/mL in a 125 mL Erlenmeyer flask containing 50 mL of CD-CHO supplemented with 4 mM glutamine. Cells were transfected with TagBFP RMCE donor and NLS-Bxb1 recombinase plasmids at a ratio of 3:1 (w/w) using FreeStyle Max transfection reagent, according to the manufacturer's instructions. Two days after transfection, cells were sub-cultured in two flasks. Three days after transfection, one flask was treated with 10 μg/mL of puromycin. Cells were passaged every three days. Eleven days after transfection, cells were subjected to flow cytometry analysis. MCL was used as the control.

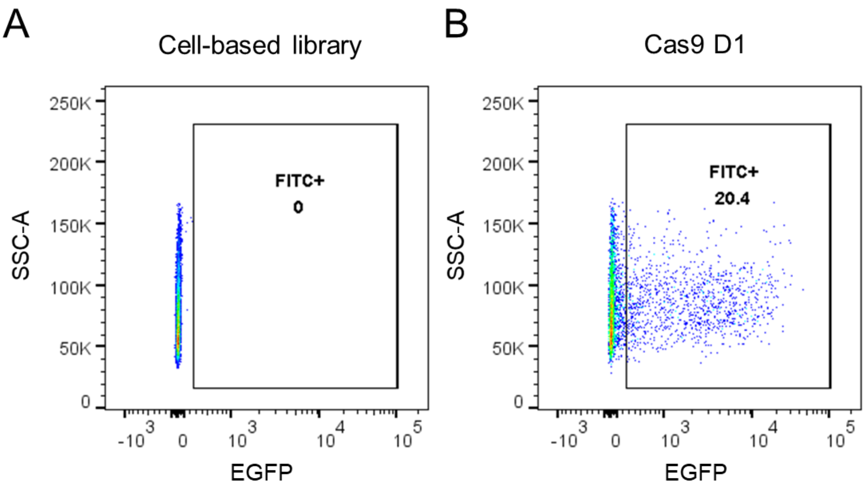


**Supplementary Fig. S3.** Cas9 transfection efficiency. Flow cytometry analysis of EGFP expression in (A) cell-based library and (B) cell-based library with Cas9 transfection. Cell-based library was seeded at a concentration of $1.0 \times 10^6$ cells/mL in a 125 mL Erlenmeyer flask containing 50 mL of CD-CHO supplemented with 4 mM glutamine. Cells were transfected with a Cas9-BSD plasmid (CMV-EGFP-T2A-Cas9-BGHpA-SV40-BSD-SV40pA) using FreeStyle Max transfection reagent, according to the manufacturer's instructions. One day after transfection, cells were subjected to flow cytometry analysis. Cell-based library was used as the control.

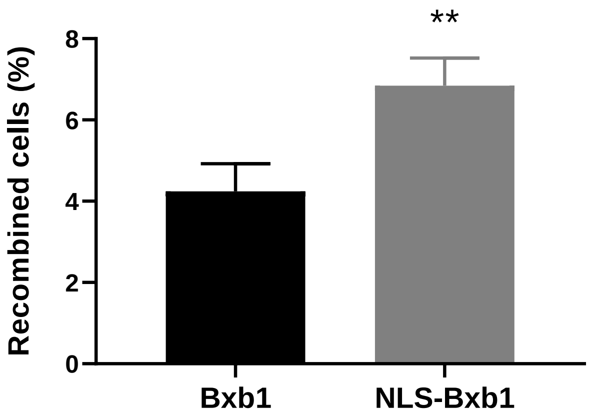


**Supplementary Fig. S4.** Determination of RMCE efficiency using Bxb1 recombinase and NLS-Bxb1 recombinase plasmids. MCL was seeded at a concentration of $1.0 \times 10^6$ cells/mL in a 6-well plate containing 3 mL of CD-CHO supplemented with 4 mM glutamine. Cells were transfected with TagBFP RMCE donor and Bxb1 or NLS-Bxb1 recombinase plasmids at a ratio of 3:1 (w/w) using FreeStyle Max transfection reagent, according to the manufacturer's instructions. Eleven days after transfection, cells were subjected to flow cytometry analysis. RMCE efficiency was measured using the percentage of mCherry-negative and TagBFP-positive cells. Asterisks (*) indicate the significant difference compared to Bxb1. Error bars in the plot represent the standard deviations of three independent experiments. An unpaired two-tailed *t*-test was used to determine the significance of the mean difference. **$P < 0.01$.

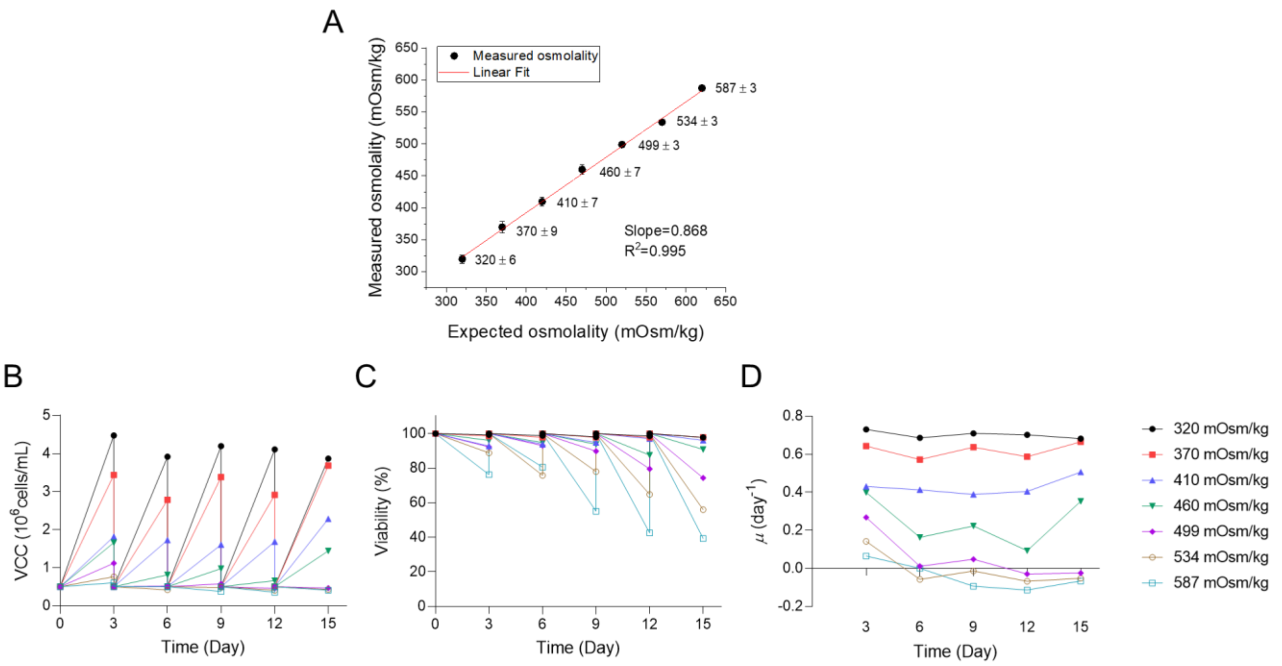


**Supplementary Fig. S5.** Effect of osmolality on cell growth. (A) Linear fit of the expected and measured osmolality. Profiles of (B) cell growth, (C) viability, and (D) specific growth rate ($\mu$) in media with various osmolality. CHO knockout library cells were seeded at a concentration of $0.5 \times 10^6$ cells/mL in 125 mL Erlenmeyer flasks containing 30 mL of CD-CHO supplemented with 4 mM glutamine and 100X anti-clumping agent with various concentrations of NaCl. To increase osmolality by 50 mOsm/kg, 150 μL of 5 M NaCl was added in 30 mL of media. NaCl was added to increase the osmolality by 0, 50, 100, 150, 200, 250, and 300 mOsm/kg. Osmolality was measured using a Fiske Micro-Osmometer. Control (320 mOsm/kg; black circle), 370 mOsm/kg (red square), 410 mOsm/kg (blue triangle), 460 mOsm/kg (green down triangle), 499 mOsm/kg (purple diamond), 534 mOsm/kg (gold empty circle), and 587 mOsm/kg (cyan empty square).

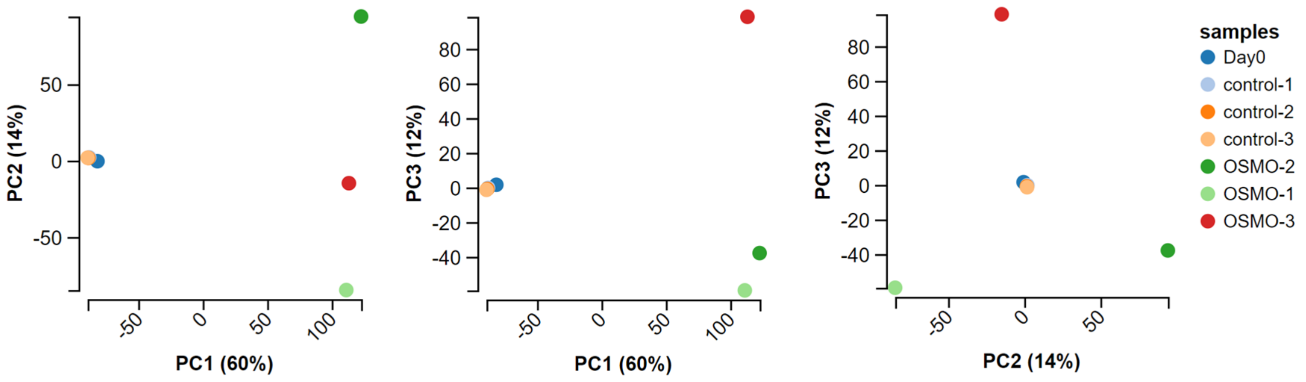


**Supplementary Fig. S6.** Principal component analysis (PCA). PCA of next-generation sequencing (NGS) data from cells sampled on day 0 (day0), cells sampled on day 21 in the standard medium (control), and cells sampled on day 21 in the hyperosmolar medium (OSMO). PCA was conducted using MAGeCK-VISPR tool (Li et al., 2015).

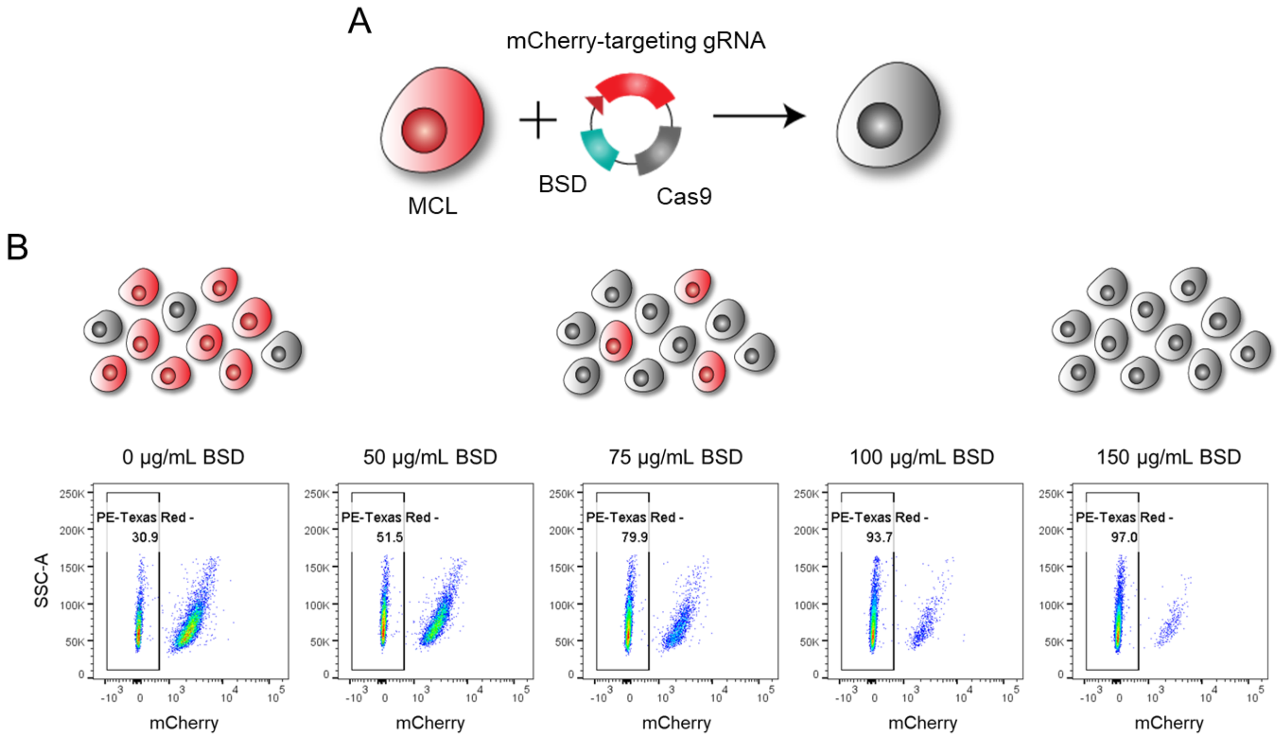


**Supplementary Fig. S7.** Measurement of mCherry knockout efficiency in CHO-K1 master cell line (MCL) using an all-in-one CRISPR/Cas9 plasmid. (A) Schematic diagram showing mCherry gene knockout. (B) Knockout efficiency after various concentrations of blasticidin selection. MCL harboring the mCherry gene was transfected with mCherry-targeting all-in-one CRISPR/Cas9 plasmid. After 48 h, transfected cells were treated with various concentrations of blasticidin for three days followed by recovery for nine days without blasticidin. Knockout efficiency of mCherry was measured using flow cytometry, as previously described (Shin et al., 2022). BSD: blasticidin.

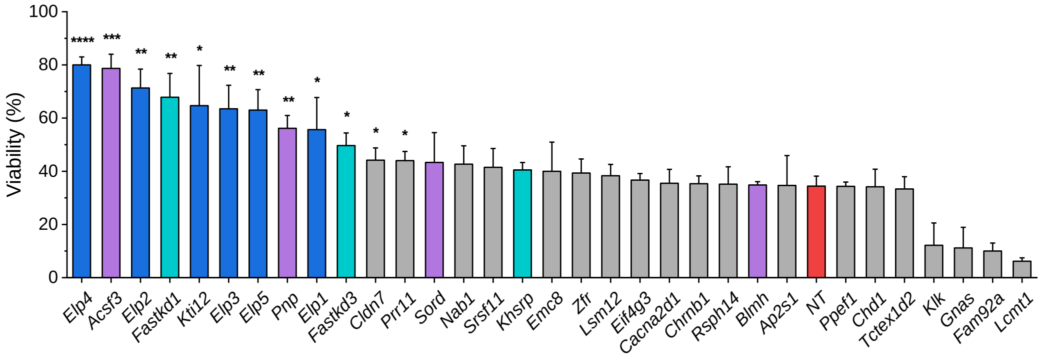


**Supplementary Fig. S8.** Viabilities of the 32 candidate gene knockout cell pools. Viability was measured on day 10 in the batch culture in the hyperosmolar medium. Columns in blue represent genes in the Gene Ontology (GO) term of tRNA wobble uridine modification, in purple represent genes in the GO term of small molecule catabolic process, in cyan represent genes in the GO term of regulation of mRNA stability and in red represent the NT control. Asterisks (*) indicate the significant difference compared to NT control. Error bars in the plot represent the standard deviations of three independent cultures. An unpaired two-tailed *t*-test was used to determine the significance of the mean difference. **P* < 0.05, ***P* < 0.01, ****P* < 0.001, and *****P* < 0.0001.

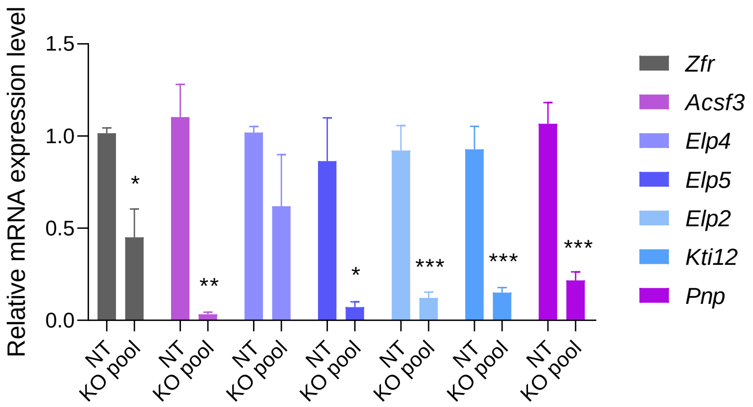


**Supplementary Fig. S9.** Relative mRNA expression levels of each target gene in CHO-mAb knockout cell pools. Exponentially growing cells were seeded at a concentration of 0.3 × $10^6$ cells/mL in 125 mL Erlenmeyer flasks containing 30 mL of CD-CHO supplemented with 4 mM glutamine and 100X anti-clumping agent. Cells were harvested on day 3. Asterisks (*) indicate the significant difference compared to the NT control. Error bars in the plot represent the standard deviations of three independent experiments. An unpaired two-tailed *t*-test was used to determine the significance of the mean difference. *$P < 0.05$, **$P < 0.01$, and ***$P < 0.001$.

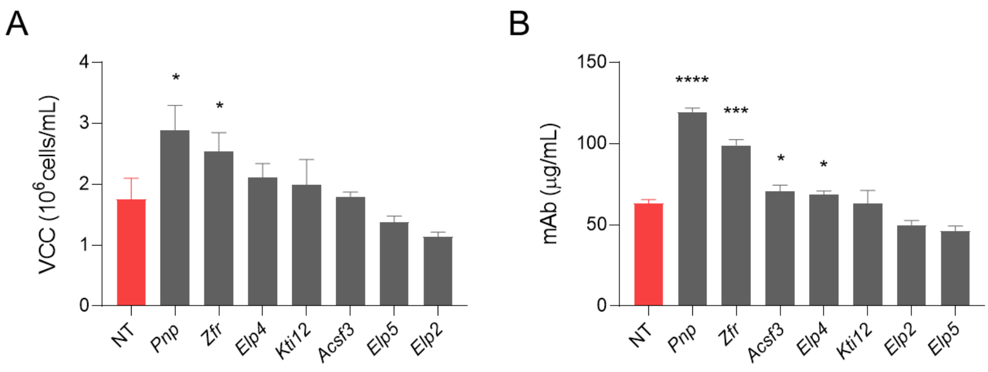


**Supplementary Fig. S10.** Assessment of gene knockout in CHO-mAb cell line. Profiles of (A) cell growth and (B) mAb concentration for CHO-mAb knockout cell pools. Exponentially growing cells were seeded at a concentration of 0.5 × $10^6$ cells/mL in 125 mL Erlenmeyer flasks containing 30 mL of CD-CHO supplemented with 4 mM glutamine, 100X anti-clumping agent, and 600 µL of 5 M NaCl. Viable cell and mAb concentrations were measured on day 6. Asterisks (*) indicate the significant difference compared to the NT control. Error bars in the plot represent the standard deviations of three independent experiments. An unpaired two-tailed *t*-test was used to determine the significance of the mean difference. *$P < 0.05$, ***$P < 0.001$, and ****$P < 0.0001$.

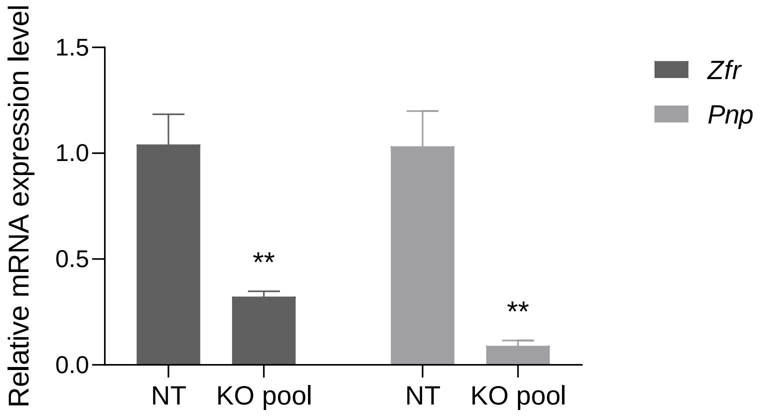


**Supplementary Fig. S11.** Relative mRNA expression levels of each target gene in CHO-bsAb knockout cell pools. Exponentially growing cells were seeded at a concentration of 0.2 × $10^6$ cells/mL in 125 mL Erlenmeyer flasks containing 30 mL of Dynamis medium supplemented with 4mM glutamine, 100 nM methotrexate, and 0.2% anti-clumping agent. Cells were harvested on day 3. Asterisks (*) indicate the significant difference compared to the NT control. Error bars in the plot represent the standard deviations of three independent experiments. An unpaired two-tailed *t*-test was used to determine the significance of the mean difference. **$P < 0.01$.

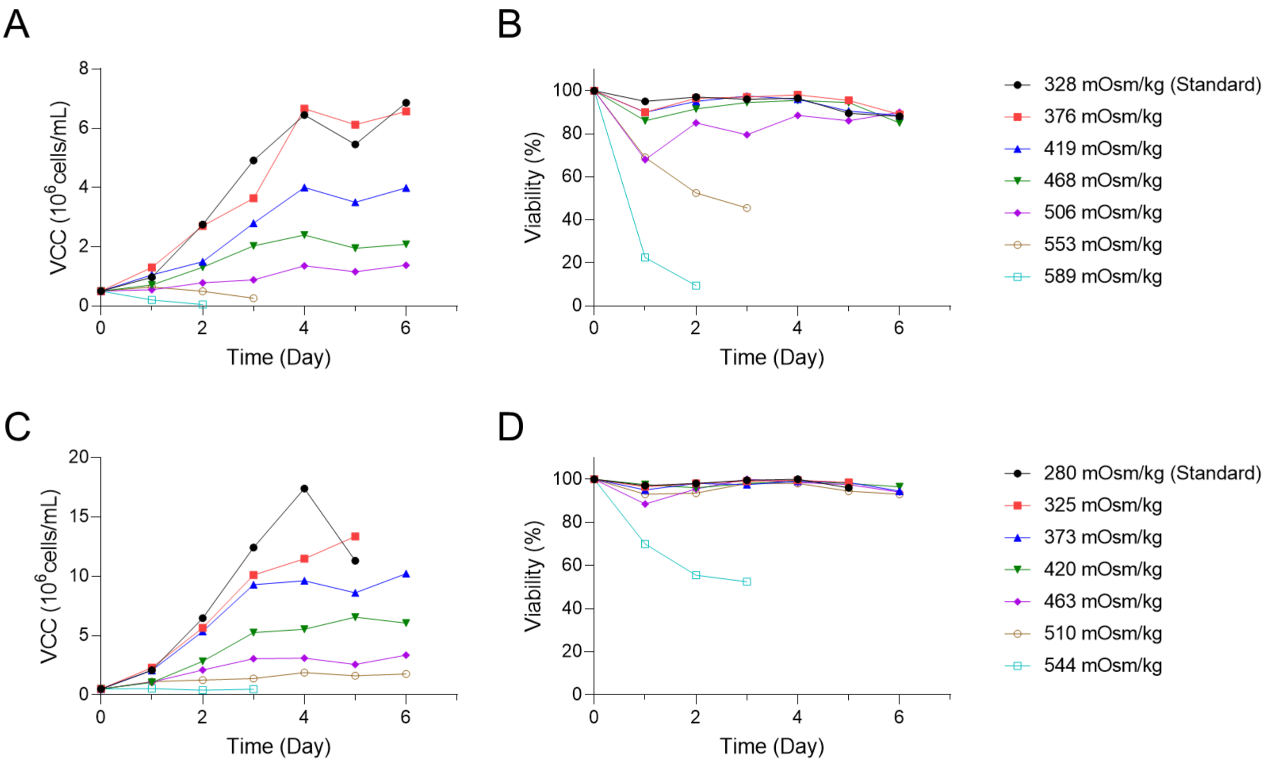


**Supplementary Fig. S12.** Effects of osmolality on CHO-mAb and CHO-bsAb cell lines. Profiles of (A) cell growth and (B) viability of CHO-mAb cell line with various osmolality. Cells were seeded at a concentration of $0.5 \times 10^6$ cells/mL in a 6-well plate containing 3 mL of PowerCHO2CD medium supplemented with GSEM and 25 µM methionine sulfoximine with various concentrations of NaCl. Profiles of (C) cell growth and (D) viability of CHO-bsAb cell line with various osmolality. Cells were seeded at a concentration of $0.5 \times 10^6$ cells/mL in a 6-well plate containing 3 mL of Dynamis medium supplemented with 4 mM glutamine, 100 nM methotrexate, and 0.2% anti-clumping agent with various concentrations of NaCl. To increase osmolality by 50 mOsm/kg, 15 µL of 5 M NaCl was added in 3 mL of media. NaCl was added to increase the osmolality by 0, 50, 100, 150, 200, 250, and 300 mOsm/kg. Osmolality was measured using a Fiske Micro-Osmometer.

**Supplementary Table S1. Plasmid information.**

| Plasmid name | Description | Reference |
|---|---|---|
| Site T2-LP donor | LP donor plasmid targeting site T2 locus (5'HA(siteT2)-EF1α-attP-mCherry-BGHpA-SV40-HygR-attB$^{mut}$-SV40pA-3'HA(siteT2)-CMV-ZsGreen1-BGHpA) | Xiong et al., (2021) |
| Site T2-gRNA | gRNA targeting Site T2 locus (U6-gRNA(siteT2)-gRNA_scaffold) | Xiong et al., (2021) |
| Cas9 | Cas9 expression plasmid (CMV-Cas9-BGHpA) | Xiong et al., (2021) |
| NLS-Bxb1 recombinase | Bxb1 recombinase with SV40_NLS sequences (CMV-NLS-Bxb1-NLS-BGHpA)- | Created in this study |
| TagBFP RMCE donor | TagBFP RMCE donor (attB-Puro-T2A-TagBFP-attB$^{mut}$) | Created in this study |
| gRNA library | gRNA library RMCE donor (attB-Puro-BGHpA-U6-gRNA(library)-gRNA_scaffold-attB$^{mut}$) | Created in this study |
| Cas9-BSD | Cas9 expression plasmid with blasticidin resistance gene (CMV-EGFP-T2A-Cas9-BGHpA-SV40-BSD-SV40pA) | Xiong et al., (2021) |
| gRNA(NT)-Cas9-T2A-BSD | All-in-one CRISPR/Cas9 plasmid non-targeting control (U6-gRNA(NT)-gRNA_scaffold-CMV-Cas9-P2A-BSD-BGHpA) | Created in this study |
| gRNA(mCherry)-Cas9-T2A-BSD | All-in-one CRISPR/Cas9 plasmid targeting mCherry sequence (U6-gRNA(mCherry)-gRNA_scaffold-CMV-Cas9-P2A-BSD-BGHpA) | Created in this study |
| gRNA(target_gene)-Cas9-T2A-BSD | All-in-one CRISPR/Cas9 plasmid targeting candidate gene (U6-gRNA(target_gene)-gRNA_scaffold-CMV-Cas9-P2A-BSD-BGHpA) | Created in this study |

LP, landing pad; HA, homology arm; NLS, nuclear localization signal; Puro, puromycin resistance gene; BSD, blasticidin resistance gene; NT, non-targeting

**Supplementary Table S2. Primer information.**

| **Primer name** | **Description** | **Sequence (5'-3')** |
|---|---|---|
| TagBFP_backbone_fwd | USER primer for TagBFP RMCE donor plasmid | AACGTGGAUACCCAGCTTTCTTGTACAAAG |
| *TagBFP_backbone_rev | USER primer for TagBFP RMCE donor plasmid | AGCAGACUTCCTCTGCCCTCTCCACTGCCGGCACCGGGCTTGCGGGTCAT |
| *TagBFP_fwd | USER primer for TagBFP RMCE donor plasmid | AGTCTGCUAACATGCGGTGACGTCGAGGAGAATCCTGGCCCAGCTAGCATGAGCGAGCTGATTAAGGAG |
| TagBFP_rev | USER primer for TagBFP RMCE donor plasmid | ATCCACGTUTTAATTAAGCTTGTGCCCCAG |
| **Bxb1_backbone_fwd | USER primer for NLS-Bxb1 recombinase plasmid | acgaaaagUtggcagttaaACACAGTCTCTGTGCCTTCTAG |
| **Bxb1_backbone_rev | USER primer for NLS-Bxb1 recombinase plasmid | ACCTTGCGCUTCTTCTTCGGggccatggtggcGGCATGACGTCTATTTCG |
| **NLS_Bxb1_fwd | USER primer for NLS-Bxb1 recombinase plasmid | AGCGCAAGGUCggaagcATGAGAGCACTGGTGGTCATCC |
| **NLS_Bxb1_rev | USER primer for NLS-Bxb1 recombinase plasmid | acttttcgUtttttcttagggctgccTGACATCCCAGTGTGCAGTC |
| NGS_fwd | Primer for NGS sample preparation | GCTTTATATATCTTGTGGAAAGGACGAAACACC |
| NGS_rev | Primer for NGS sample preparation | CCGACTCGGTGCCACTTTTTCAA |
| Zfr_fwd | *Zfr* primer for qRT-PCR | CGAGAAGAGAACATGAGGGAAG |
| Zfr_rev | *Zfr* primer for qRT-PCR | GTTGACAAAGGTCTCGGAGAATA |
| Acsf3_fwd | *Acsf3* primer for qRT-PCR | GAACTGGAAGGAGTAGGTCTTG |
| Acsf3_rev | *Acsf3* primer for qRT-PCR | GCCAGCAGTTCTGGTATCTT |
| Elp4_fwd | *Elp4* primer for qRT-PCR | CTTCTATGTTCTCCGTGGTCTTC |
| Elp4_rev | *Elp4* primer for qRT-PCR | ACACGAGCAGTAATTGCTTTATTC |
| Elp5_fwd | *Elp5* primer for qRT-PCR | CCAGACAACTAGCCACATCTT |
| Elp5_rev | *Elp5* primer for qRT-PCR | GCAGTCTTCCTCTCCCATTT |
| Elp2_fwd | *Elp2* primer for qRT-PCR | GTCACTGGATCACTTGCTTTATG |
| Elp2_rev | *Elp2* primer for qRT-PCR | CTCCCGGATAGACAGCTTTATT |
| Kti12_fwd | *Kti12* primer for qRT-PCR | GGAACTGGATCCGGAAGAAAT |

| | | |
|---|---|---|
| Kti12_rev | *Kti12* primer for qRT-PCR | GGAGGAAAGGCACTGGAAG |
| Pnp_fwd | *Pnp* primer for qRT-PCR | ACATCAACCTACCTGGTTTCTC |
| Pnp_rev | *Pnp* primer for qRT-PCR | CCGGTCATAAGCATCAGACAT |
| Gapdh_fwd | *Gapdh* primer for qRT-PCR | GGACATCAAGAAGGTGGTGAA |
| Gapdh_rev | *Gapdh* primer for qRT-PCR | GAGTGGGAGTCACTGTTGAAG |

fwd, forward primer; rev, reverse primer

*T2A sequences are included in the primers.

**Nuclear localization signal (NLS) sequences are included in the primers.

**Supplementary Table S3. gRNA information.**

| **Target** | **gRNA sequence (N20NGG)** |
|---|---|
| Site T2 locus | GAGATCTGTGGGCCGAGTAAGGG |
| Non targeting | ACACCGTTCCGAGTTCGGTT |
| mCherry | GGCCACGAGTTCGAGATCGAGGG |
| *Fastkd1* | GCTGACCATCAAGACAGACAGGG |
| *Zfr* | TATGTACTGGAGACGAATGGAGG |
| *Acsf3* | ACTCTACTGGAAGCACCCTGAGG |
| *Elp4* | ACGGAAGCTGTTCACCATTGAGG |
| *Elp5* | TTAAAGGTCAAATCAGTCGTGGG |
| *Lcmt1* | CATCATGGAGTTACACCCAGAGG |
| *Eif4g3* | GTTCAAAGAGTGGATGAAGGTGG |
| *Elp2* | AGACTTAATATGGGATCCAGAGG |
| *Gnas* | CTACAACATGGTCATTCGGGAGG |
| *Kti12* | GGACTCGGTGAACTACATCAAGG |
| *Elp1* | GGGCTCGGAAGATCAGAGTGTGG |
| *Fastkd3* | AGAAGCTTCAGTAGCAACTGTGG |
| *Cacna2d1* | TCACGAAATCATCATCCGAGAGG |
| *Ppef1* | TATTCAGGTGCAGGTCAGTGGGG |
| *Blmh* | CCCTGAATCTCATACGACAGAGG |
| *Khsrp* | GTAGTAATGGGAGTAGTAGGCGG |
| *Cldn7* | TAACATTCATGGGTGTCAAGGGG |
| *Elp3* | CCCAGTTCCACGGATCACCAGGG |

| | |
|---|---|
| *Emc8* | CGCCAAGTACCCGCACTGCG<u>CGG</u> |
| *Srsf11* | GAATGCTTACTTGGGATCAA<u>CGG</u> |
| *Chrnb1* | AAACACAGTCAGGGTCAGAA<u>GGG</u> |
| *Rsph14* | TAATGAGGAGCTGCAATCGG<u>AGG</u> |
| *Sord* | TGCTTGGGCATGAAGCTGCA<u>GGG</u> |
| *Prr11* | CTCCACCACCCACGCCAGAA<u>AGG</u> |
| *Ap2s1* | GTAGATGAGATGTTCCTGGC<u>AGG</u> |
| *Fam92a* | TCAAAGCAACATTAACAGCA<u>AGG</u> |
| *Pnp* | GTGAGTATCCTTCATATGTG<u>TGG</u> |
| *Klk7* | ACAGTGCAGGATGTCCCTGG<u>GGG</u> |
| *Nab1* | AGTACACTTACAGGACCACC<u>AGG</u> |
| *Lsm12* | CCAGAATGTCCCTCTTCCAG<u>TGG</u> |
| *Chd1* | TTAATTCGCCTAAGAGAAAG<u>AGG</u> |
| *Tctex1d2* | ACATCTGTCAGAAACCATTA<u>AGG</u> |
